# CXCL8 secreted by immature granulocytes inhibits wildtype hematopoiesis in chronic myelomonocytic leukemia

**DOI:** 10.1101/2024.03.08.583935

**Authors:** Paul Deschamps, Margaux Wacheux, Axel Gosseye, Margot Morabito, Arnaud Pagès, Anne-Marie Lyne, Alexia Alfaro, Philippe Rameau, Aygun Imanci, Rabie Chelbie, Valentine Marchand, Aline Renneville, Mrinal Patnaik, Valerie Lapierre, Bouchra Badaoui, Orianne Wagner-Ballon, Céline Berthon, Thorsten Braun, Christophe Willekens, Raphael Itzykson, Pierre Fenaux, Sylvain Thépot, Gabriel Etienne, Francoise Porteu, Emilie Elvira-Matelot, Nathalie Droin, Leïla Perié, Lucie Laplane, Eric Solary, Dorothée Selimoglu-Buet

## Abstract

Chronic myelomonocytic leukemia (CMML) is a severe myeloid malignancy with limited therapeutic options. Single-cell analysis of clonal architecture demonstrated early clonal dominance with few residual wildtype hematopoietic stem cells. Circulating myeloid cells of the leukemic clone and the cytokines they produce generate a deleterious inflammatory climate. Our hypothesis is that therapeutic control of the inflammatory component in CMML could contribute to stepping down disease progression. The present study explores the contribution of immature granulocytes (iGRANs) to CMML progression. iGRANs can be detected and quantified in the peripheral blood of patients by spectral and conventional flow cytometry. Their accumulation is a potent and independent poor prognostic factor. These cells belong to the leukemic clone and behave as myeloid-derived suppressor cells. Bulk and single cell RNA sequencing revealed a pro-inflammatory status of iGRAN that secrete multiple cytokines of which CXCL8 at the highest level. This cytokine inhibits the proliferation of wildtype but not CMML hematopoietic stem and progenitor cells (HSPCs) in which CXCL8 receptors are epigenetically downregulated. CXCL8 receptor inhibitors and CXCL8 blockade restore wildtype HSPC proliferation, suggesting that relieving CXCL8 selective pressure on wildtype HSPCs is a potential strategy to slow CMML progression and restore some healthy hematopoiesis.

## Introduction

Chronic myelomonocytic leukemia (CMML) is a myeloid malignancy defined by overlapping features of both myeloproliferative and myelodysplastic neoplasms (1). CMML diagnosis is based on sustained absolute and relative peripheral blood monocytosis with abnormal partitioning of peripheral blood monocyte subsets, one of more clonal cytogenetic or molecular abnormality, and/or dysplasia in at least one lineage (2). The disease predominantly affects elderly patients and mutational signatures of leukemic cells suggest that ageing is the main cause of the disease (3). Multiple studies have substantiated the clinical and molecular distinction of dysplastic (MD-CMML) and proliferative (MP-CMML) CMML on the basis of cutoff white blood cell (WBC) count of 13 x 10^9^/L (4). Blast cell count in the bone marrow also separates CMML in subgroups with distinct outcome (5). A watch and wait attitude with careful monitoring is proposed for lower risk patients while severe CMML requires therapy (1). Allogeneic stem cell transplantation, which is the only potentially curative treatment, is commonly precluded by age and comorbidities, and only partially abrogates the risk of relapse (6). In patients with MD-CMML, hypomethylating agents can restore balanced hematopoiesis but do not reduce the variant allele frequency among circulating myeloid cells (3) and do not prevent progression to acute myeloid leukemia (1). In patients with MP-CMML, hypomethylating agents do not demonstrate a survival benefit compared to cytoreductive therapy with hydroxyurea (7). Therefore, there is an urgent need for additional therapeutic approaches for this disease.

CMML is a clonal disorder driven by the linear accumulation, in the hematopoietic stem cell (HSC) compartment, of somatic variants that diversely affect DNA methylation, histone modifications, pre-mRNA splicing and cell signaling (4). Clonal architecture analyzed at the single cell level indicates early clonal dominance with a very low number of residual wildtype HSCs in the bone marrow (8). Myeloid differentiation of mutated HSCs is amplified by hypersensitivity of myeloid progenitor cells to granulocyte/macrophage colony-stimulating factor (GM-CSF) (9). Virtually all the mature myeloid cells circulating in the body belong to the malignant clone (10,11).

While 15-30% of CMML patients die from disease transformation into acute leukemia, most of them demonstrate insidious physical exhaustion in an inflammatory climate. Circulating myeloid cells, predominantly monocytes and neutrophils, may contribute to the elevated levels of pro-inflammatory cytokines detected in the plasma of CMML patients (12,13). Analysis of gene expression in peripheral blood monocytes indicates a pro-inflammatory phenotype (10,14). In mouse models of chronic myeloid malignancies, inflammatory cytokines secreted by mature myeloid cells of the leukemic clone promote HSCs expansion (15,16). Therapeutic inhibition of these cytokines prevents disease development and progression (16,17). Such feed-forward loops involving inflammatory cytokines produced by myeloid cells of the leukemic clone could contribute also to the progression of CMML, *e.g.*, by promoting the expansion of mutated HSCs or slowing down that of wildtype cells.

Looking for the respective contribution of myeloid cell subsets of the leukemic clone to the inflammatory climate observed in CMML, we focused the present study on the granulocytic lineage. Neutrophil precursors, including metamyelocytes, myelocytes and promyelocytes, which are normally retained in the bone marrow, are detected cytologically in the peripheral blood of a fraction of CMML patients and referred to as immature myeloid cells (IMC) (13,18). We used single cell approaches to define immature granulocytes and analyze further their contribution to disease outcome and pathogenesis. In two independent cohorts of patients, the presence of immature granulocytes in the peripheral blood correlates with a poor outcome. These cells, which share the same clonal origin as monocytes, demonstrate an inflammatory and immunosuppressive phenotype. They secrete high levels of CXCL8, a cytokine that inhibits the proliferation and differentiation of wildtype HSCs while sparing mutated HSCs in which CXCL8 receptors are downregulated as consequence of clonal epigenetic changes. Reparixin, a clinical drug targeting the CXCL8-CXCR1/2 axis as well as CXCL8 neutralizing antibody, restore the proliferation of wildtype HSCs. The feed-forward loop that involves CXCL8 produced by dysplastic granulocytes of CMML leukemic clone in the repression of residual wildtype HSC suggests a new strategy for the therapeutic management of CMML patients.

## Results

### Identification and characterization of immature granulocytes

We first assessed the prognostic significance of IMC detection ≥1% on routine blood smears in a cohort of 580 consecutive CMML patients at diagnosis from Mayo Clinic (median age: 71 years [18-95]; 68% males) (**Supp Table 1**). The overall survival (OS) of the 351 patients with IMC was significantly lower than that of the 229 patients without IMC, suggesting a poor prognostic factor. However, when WHO-defined criteria used for CMML stratification, namely WBC count and blast cell fraction, were considered together with IMC subgroups in multivariate analysis, IMC and blast cell count were no longer significant (**Supp Figure 1A**). Actually, IMC is not part of most scoring systems proposed in CMML, which may be related to the limited reproducibility of IMC quantification on routine complete blood counts (19). We explored whether flow cytometry phenotyping of peripheral blood cells could refine IMC identification and characterization.

**Figure 1.**
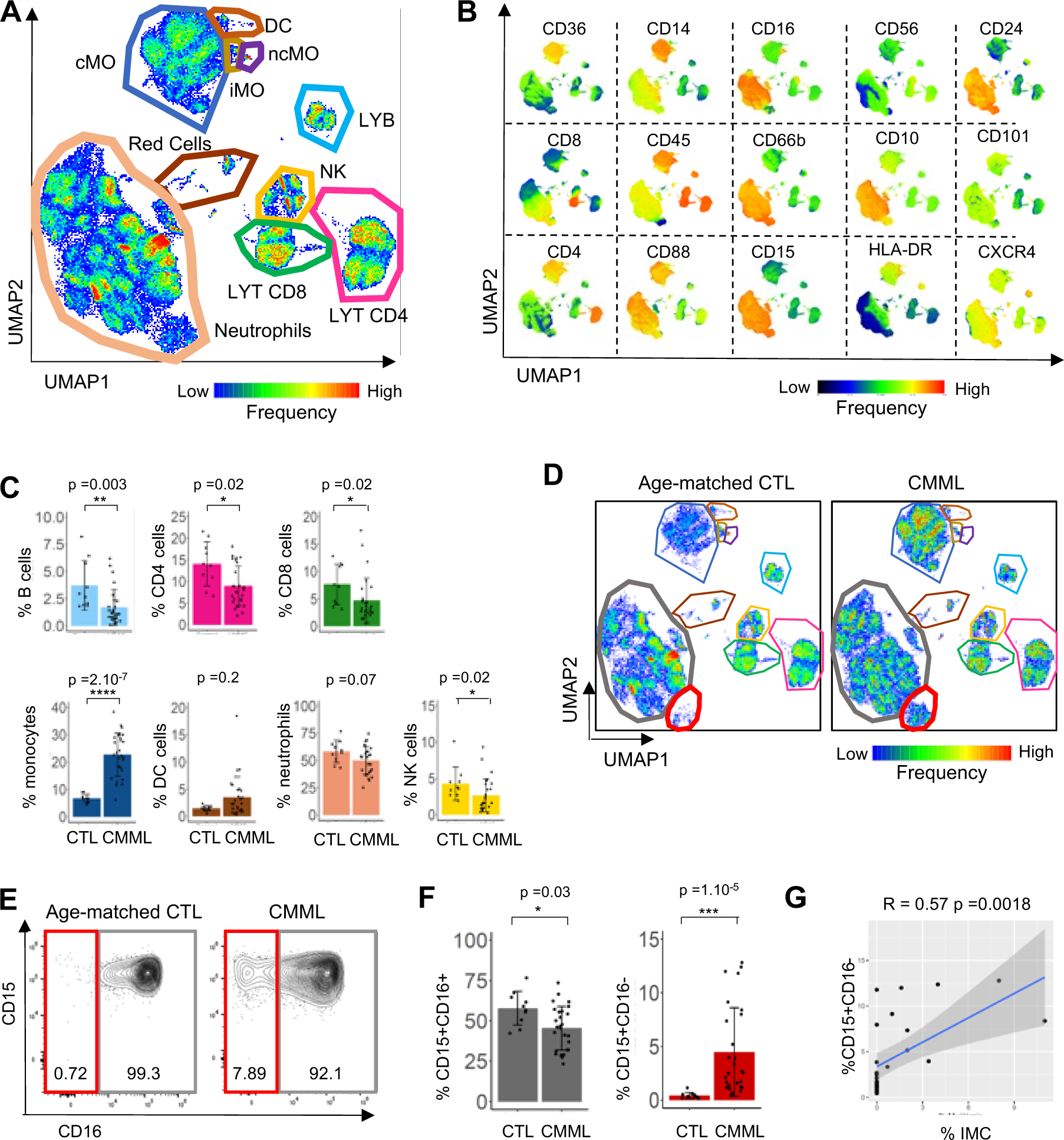
Spectral flow analysis identifies immature granulocytes (iGRANs) in CMML peripheral blood. **A.** Non-supervised UMAP of spectral flow cytometry analysis of peripheral blood cells collected from 27 CMML patients and 10 age-matched control donors. cMO : classical monocytes, iMO : intermediate monocytes, ncMO : non-classical monocytes, LYB : B lymphocytes, LYT : T lymphocytes, DC : Dendritic cells, NK : Natural killer cells. **B**. Cell surface expression of indicated markers on the UMAP shown in A. **C**. Fraction of B, CD4^+^ T, CD8^+^ T, monocytes, DC, neutrophils and NK cells among total CD45^+^ cells in age-matched controls (CTL) and CMML patients. Mann-Whitney test. **D**. Non-supervised UMAP analysis of spectral flow cytometry data in the 10 controls compared to 27 CMML patients (60,000 cells for each condition). **E**. Partition of neutrophil subsets based on CD15 and CD16 expression, in each group. **F**. Percentage of CD15^+^,CD16^-^ and CD15^+^,CD16^+^ neutrophils as separated in E, among CD45^+^ cells. **G.** Spearman correlation between CD15^+^CD16^-^ and immature myeloid cell (IMC) fractions in CMML peripheral blood. Adjusted p-values are indicated above the graphs.

We used spectral flow cytometry with a panel of 34 antibodies recognizing cell surface markers (**Supp Table 2**) to generate an overview of cell populations in the peripheral blood. Pooled data from untreated CMML patients (N=27) and age-matched controls (N=10) (**Supp Table 3**) were subjected to a clustering analysis and visualized after a dimensionality reduction using the unsupervised uniform manifold approximation and projection (UMAP) algorithm (20) (**Figure 1A, Supp Figure 1B).** This approach allowed identifying classical, intermediate and nonclassical monocytes; dendritic cells; B cells; CD4^+^ and CD8^+^ T cells; natural killer (NK) cells; and neutrophils (**Figure 1B).** A gating strategy was applied to precisely quantify each cell population in every sample among white blood cells (CD45^+^ cells, **Supp Figure 1C-J**). Examination of the cell-type profiles of samples from patients with CMML and controls revealed the increased fraction of circulating monocytes that defines CMML (**Figure 1C**), with a typical accumulation of classical monocytes at the expense of intermediate and nonclassical monocytes (**Supp Figure 1K)**. The fractions of circulating B cells, CD4^+^ and CD8^+^ T cells and NK cells decreased in CMML patients compared to age-matched controls, whereas no significant changes were observed in the global dendritic cell and granulocyte populations (**Figure 1C**).

A detailed analysis of neutrophils detected a cluster of cells in CMML samples that was hardly observed in age-matched controls (**Figure 1D**). This cell subset expressed high levels of CD15, CD24 and CXCR4; low levels of CD45, CD10, and CD101; and almost no CD16, suggestive of immature granulocytes (iGRANs) (**Figure 1B**). Quantification of the neutrophil populations among CD45^+^ cells in individual samples revealed a significant decrease in the fraction of mature CD15^+^CD16^+^ neutrophils in the CMML group compared to the healthy donor group, while the fraction of CD15^+^CD16^-^ cells significantly increased with high interpatient heterogeneity (**Figure 1E, 1F**). In the CMML group, the fraction of CD15^+^CD16^-^ neutrophils correlated with the IMC fraction detected on routine blood smears (r=0.57, **Figure 1G**). Altogether, spectral flow cytometry analysis of CMML peripheral blood cells refined the detection of IMC and characterized a subset of cells as CD15^+^CD16^-^CD66b^+^ iGRANs.

### Flow cytometric detection of iGRANs refines CMML prognostication

Having identified informative phenotypic markers by spectral flow cytometry analysis, we developed a conventional, multiparametric flow cytometry assay (**Supp Table 2**) to routinely quantify the fraction of iGRANs in freshly collected blood samples and revisit the prognostic significance of this parameter in CMML. We noticed that iGRANs were part of the peripheral blood mononucleated cell (PBMC) population sorted by low density gradient centrifugation, a process that removes a majority of CD15^+^CD16^+^CD66b^+^ mature neutrophils without affecting iGRAN population (**Supp Figure 2A, 2B**). Based on this low-density cell property, the conventional flow cytometry assay was performed on PBMC (**Supp Figure 2C-J**). The iGRAN fraction measured among Lin^-^CD16^-^CD11b^+^CD33^+^ cells correlated with that among total CD45^+^ cells (**Supp Figure 2K**). With this conventional flow cytometry assay, repeated measures of the iGRAN fraction over up to one year in 17 untreated CMML patients showed reproducible results (**Supp Figure 2L**).

**Figure 2.**
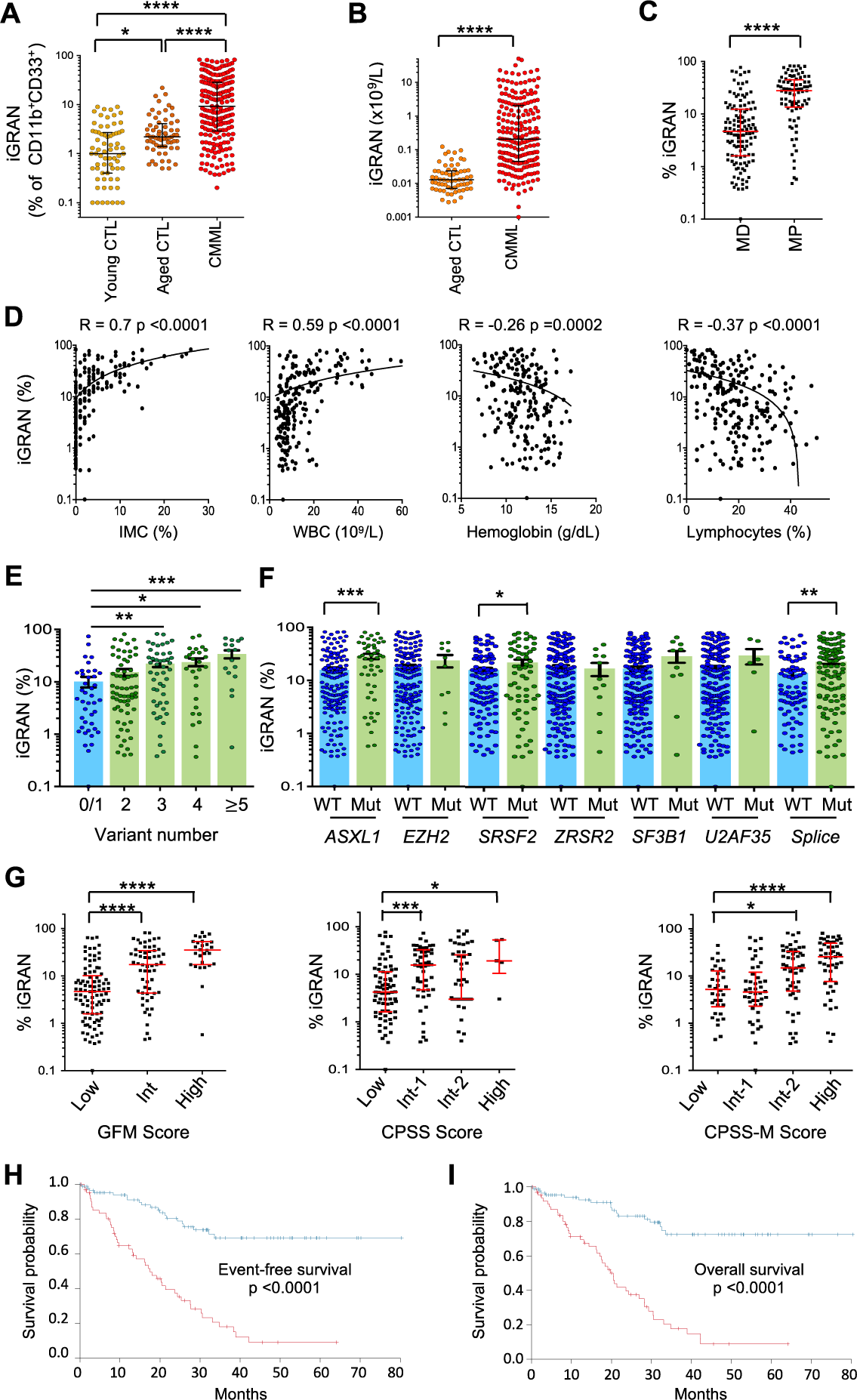
Elevated iGRAN fraction in the peripheral blood of CMML patients is a poor prognostic factor. **A**. iGRAN fraction in CD11b^+^CD33^+^ population as measured by conventional flow cytometry in the peripheral blood of young controls (N=71), age-matched controls (N=64) and CMML patients (N=209). Kruskal-Wallis non-parametric test. **B**. iGRAN absolute number (×10^9^/ L) was measured in the peripheral blood of CMML patients compared to age-matched controls. **C**. iGRAN fraction in dysplastic (MD) and proliferative (MP) CMML subtypes according to the WHO classification. **D**. Spearman correlation between iGRAN fraction and immature myeloid cell (IMC) fraction, white blood cell count (WBC), hemoglobin level and lymphocyte fraction in the peripheral blood of CMML patients. **E**. iGRAN fraction in CMML patients grouped according to the number of mutations detected in a panel of 25 genes. Kruskal-Wallis non-parametric test. **F**. iGRAN fraction in CMML patients grouped according to the mutational status of each indicated gene: wildtype (WT) or mutated (Mut). *Splice*: *SRSF2+ZRSR2+U2AF1+SF3B1*. **G**. iGRAN fraction in CMML patients grouped according to GFM, CPSS and CPSS-Molecular (CPSS-M) prognostic scores. Kruskal-Wallis non-parametric test. **H,I**. Event-free survival (EFS, defined as time between diagnosis and AML transformation, death, or last follow-up) (H) and overall survival (time between diagnosis and death)(I) of CMML patients with high (≥14%, N=66, in red) or low (<14%, N=88, in blue) iGRAN fraction; log-rank test.

Once calibrated, this flow cytometry assay was prospectively assessed between March 2015 and April 2019 on a learning cohort of 209 untreated CMML patients (median age: 75 years [50-93], 63% males) compared to 64 age-matched controls (median age: 74 years [65-94]) and 71 younger healthy donors (< 65 years old) (**Supp Table 3**). CMML diagnosis was supported by flow cytometry analysis of peripheral monocyte subsets (21): in 192 patients (92%), classical monocytes represented more than 94% of total monocytes, while a decreased fraction of slan^+^ nonclassical monocytes was observed in 15 of the 17 remaining patients (7%)(22,23). The fraction of iGRANs among myeloid cells was significantly higher in aged controls (median 2.2% [0.5-22%]) than in younger ones (median 1.0% [0-0.9%], p=0.0001) and was further increased in CMML patients (median 9.2% [0.03-82.5%], p<0.0001) (**Figure 2A**). The absolute number of circulating iGRANs also was statistically increased in CMML patients, sometimes reaching values close to 50×10^9^/L, representing a 1000-fold increase over the values in aged controls (**Figure 2B**). The fraction of iGRANs was significantly higher in MP-CMML than in MD-CMML (**Figure 2C**) and positively correlated with the fraction of IMCs quantified on blood smears (r=0.7, p<0.0001). However, the flow cytometric quantification of iGRANs has a more precise detection threshold for patients with low IMC (**Figure 2D**). The fraction of iGRANs also correlated with WBC count and with lower hemoglobin level and lymphocyte fraction (**Figure 2D**). In contrast, no correlation was observed for iGRAN fraction with age, sex, presence of cytogenetic abnormalities or total neutrophil fraction, and the correlations with monocyte fraction, platelet count and WHO-defined subtypes remained weak (**Supp Figure 3 A-E**). Analysis of mutations in a panel of 35 genes recurrently mutated in CMML (**Supp Table 3**) showed an increase in the iGRAN fraction with an increasing number of mutated genes as well as with the presence of *ASXL1* or *SRSF2* gene mutations (**Figure 2E-F**). No significant association was detected with the other common variants in CMML, including those in signaling genes (**Supp Figure 3F**). Finally, the iGRAN fraction increased with disease risk level, as measured using the Groupe Francophone des Myélodysplasies score (GFM)(24), the CMML-specific prognostic scoring system (CPSS)(25) and the CPSS-M score that incorporates molecular genetic data (26)(**Figure 2G**). Together, these results indicate that iGRAN accumulation in blood is associated to poor prognosis scores and to poor prognosis markers like *ASXL1* mutations (27), or proliferative forms of the disease. The relationship between the iGRAN fraction and other parameters were validated by analyses performed using the absolute number of circulating cells (10^9^/L) (**Supp Table 4**), indicating the robustness of iGRAN quantification to separate the most severe forms of the disease.

**Figure 3.**
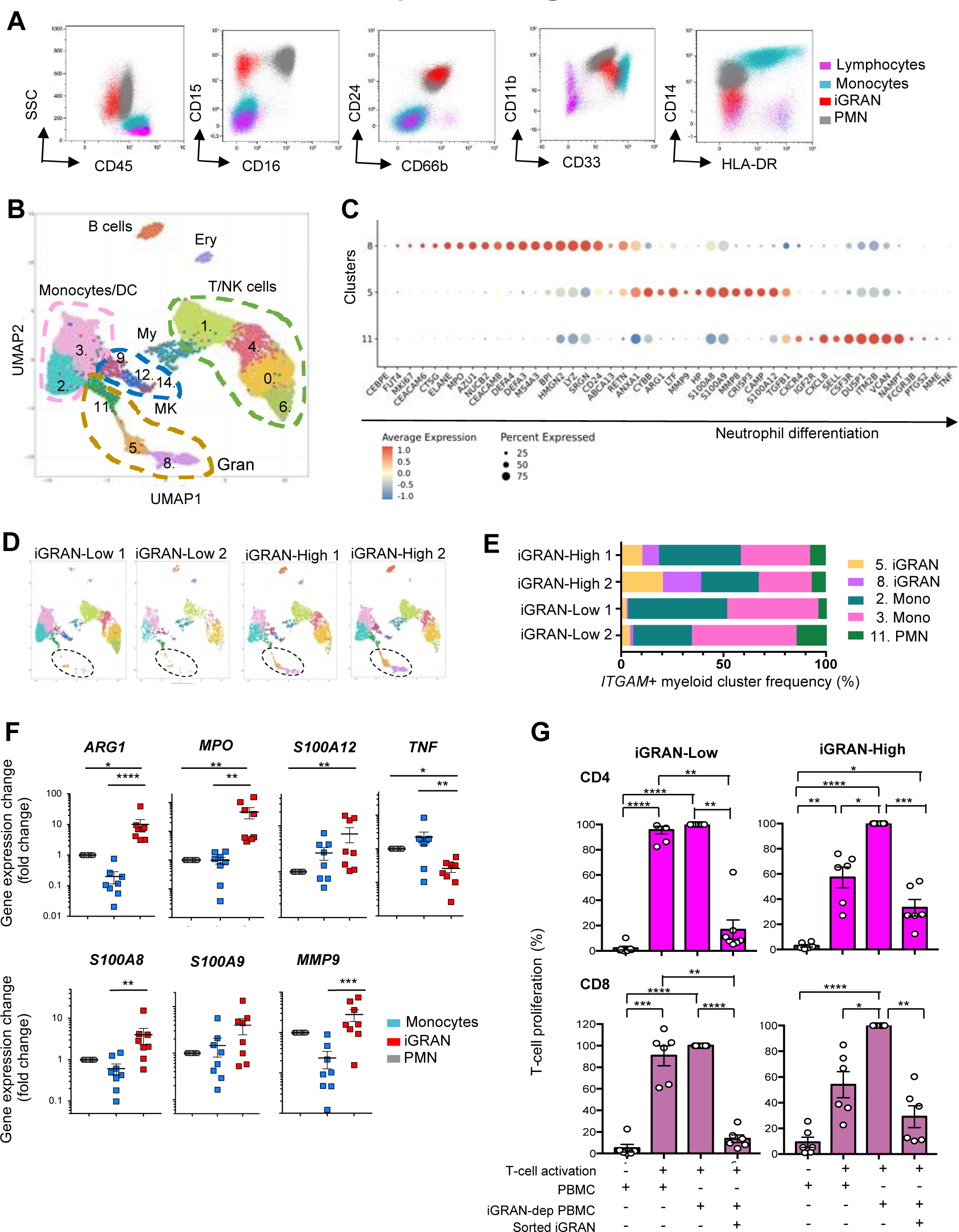
iGRANs demonstrate features of myeloid-derived suppressive cells. **A**. Representative conventional flow plots separating iGRANs (CD45^low^, CD33^+^, CD11b^+^, HLA-DR^-^, CD14^-^, CD15^+^, CD24^+^, CD66b^+^ cells) from monocytes, PMN and lymphocytes in low density peripheral blood cells (PBMC). **B**. Single cell analysis of PBMC collected from two CMML patients with a fraction ≥14% and two with a lower iGRAN fraction; unsupervised clustering of pooled data separating 15 clusters groups in indicated cell categories (Ery, erythroid cells; DC, dendritic cells; My, myeloid cells; MK, megakaryocytes; Gran, granulocytes). **C**. Dot plot showing the average expression (color-scaled) of selected granulocyte genes and the percentage of cells that expressed those genes in indicated granulocytic clusters. **D**. UMAP representation of each patient sample, two iGRAN-Low (<14%) and two iGRAN-High (≥14%) CMML patients. **E**. Fraction of each cell type in ITGAM^+^ clusters corresponding to CD11b^+^CD33^+^ cells per CMML patient. Colors are cluster codes defined in B. **F**. Expression of indicated genes in enriched fraction of iGRANs, monocytes and neutrophils measured by RT-qPCR and normalized to *RPL32*, *GUS* and *GAPDH* housekeeping genes (N=8 CMML patients). **G.** Suppressive activity of iGRANs on T cell proliferation. iGRAN-Low (<14%, N=6, left panels) and iGRAN-High (≥14%, N=6, right panels) PBMCs were labeled with Cell trace Violet before activating T-cells with anti-CD3 and anti-CD28 antibodies; CD4 (upper panels) and CD8 (lower panels) T-cell proliferation was measured at day 4 by flow cytometry. We used PBMCs without any manipulation (PBMCs), PBMCs in which iGRAN have been depleted (iGRAN-dep PBMC) and iGRAN-dep PBMC with addition of 10% sorted iGRANs. T-cell proliferation (%) is relative to the highest proliferation, observed with iGRAN-dep PBMCs.

Among the 209 patients included in the learning cohort, iGRAN quantification had been performed for 154 patients at diagnosis, *i.e.*, ± 6 months from initial bone marrow examination. With a median follow-up of 34 months, 27 patients among 154 progressed to AML, and 62 died. The median OS and median event-free survival (EFS, defined as the time between diagnosis and AML transformation, death, or last follow-up) were 20.6 and 20 months, respectively. Univariate and multivariate Cox models were built with continuous iGRAN percentages and absolute numbers. The multivariate analysis model included clinical and mutation variables with independent prognostic value, based on the GFM scoring system that includes age, WBC, hemoglobin level, platelet count and *ASXL1* mutation (24). Both univariate and multivariate models showed a significant impact of iGRANs measured at diagnosis on EFS or OS (**Supp Table 5**). We then computed the data to dichotomize continuous variables by selecting cutoff values to maximize the log-rank statistics and picked out an iGRAN fraction ≥14% of circulating myeloid cells or an absolute iGRAN number ≥0.4×10^9^/L as optimal values to identify the impact of iGRANs on EFS and OS (**Figure 2H-I** and **Supp Figure 3G**). With these cutoff values, the iGRAN fraction improved the accuracy of existing prognostic scores, after adjusting for potential confounders (hazard ratio for iGRAN fraction in multivariate model: 3.678, 95% CI: [1.850–7.305] for OS and 3.003, 95% CI: [1.592–5.663] for EFS) (**Supp Table 5**). An increased number and fraction of iGRAN was also detected in a validation cohort of 160 CMML patients (**Supp Table 3** and **Supp Figure 3H**), especially those with a proliferative phenotype (**Supp Figure 3I**), correlating again with IMC fraction, WBC, hemoglobin level and lymphocyte fraction (**Supp Figure 3J**). Follow-up (median, 36.6 months) was available for 110 of these patients, of whom 13 progressed to AML and 43 died. Using cut-off values defined within the learning cohort (iGRAN fraction ≥14%, iGRAN number ≥0.4×10^9^/L), the impact of iGRANs on EFS and OS was validated (**Supp Figure 3K**).

Together, flow cytometric measurement of an iGRAN fraction ≥14% of circulating myeloid cells, or an absolute number of iGRAN ≥0.4×10^9^/L, is an independent biomarker of CMML severity.

### iGRANs are myeloid-derived suppressive cells

Having identified the poor prognostic significance of iGRAN excess, we next sought to assess their functional impact. The phenotype of these cells (CD45^low^, CD33^+^, CD11b^+^, HLA-DR^-^, CD14^-^, CD15^+^, CD24^+^, CD66b^+^, and low density) suggested that they may be granulocytic myeloid-derived suppressor cells (G-MDSCs, also referred to as PMN-MDSCs) (**Figure 3A**)(28). To explore this possibility, we selected two patients with an iGRAN-high (≥14%) and two patients with an iGRAN-low fraction and performed single-cell RNA sequencing analysis (scRNAseq) of PBMCs. The data were pooled and subjected to dimensionality reduction using the UMAP algorithm, and 15 clusters were identified (**Figure 3B**). Based on the expression of marker genes, the cell types of clusters were identified, including T/NK cells (*CD3E, NKG7*), B cells (*CD79*A), erythroid cells (*HBA1*), megakaryocytes/platelets (*PF4*), monocytes (*CD14, CD33*), and granulocytes (*FCGR3B, S100A9, S100A8,* and *LYZ*) (**Supp Figure 4A-C**). Granulocytes encompassed several clusters (11, 5, and 8), corresponding to sequential stages of neutrophil maturation. Cells in cluster 8 expressed genes encoding primary neutrophil granule proteins, such as *ELANE, CEACAM, AZU1, DEFA1* and *DEFA3*, indicating promyelocytes and myelocytes (**Figure 3C, Supp Figure 4B**). Cells in cluster 5 expressed genes encoding secondary and tertiary neutrophil granule proteins, such as *MMP9, MMP8* and *LTF*, indicating metamyelocytes. Lastly, cells in cluster 11 expressed high levels of genes that characterize mature neutrophils, such as *MME* (CD10) and *FCGR3B* (CD16). Patients with an iGRAN-high fraction showed strong enrichment of cells in clusters 5 and 8 (**Figure 3D-E**), validating the initial description of this population as immature granulocytes.

**Figure 4.**
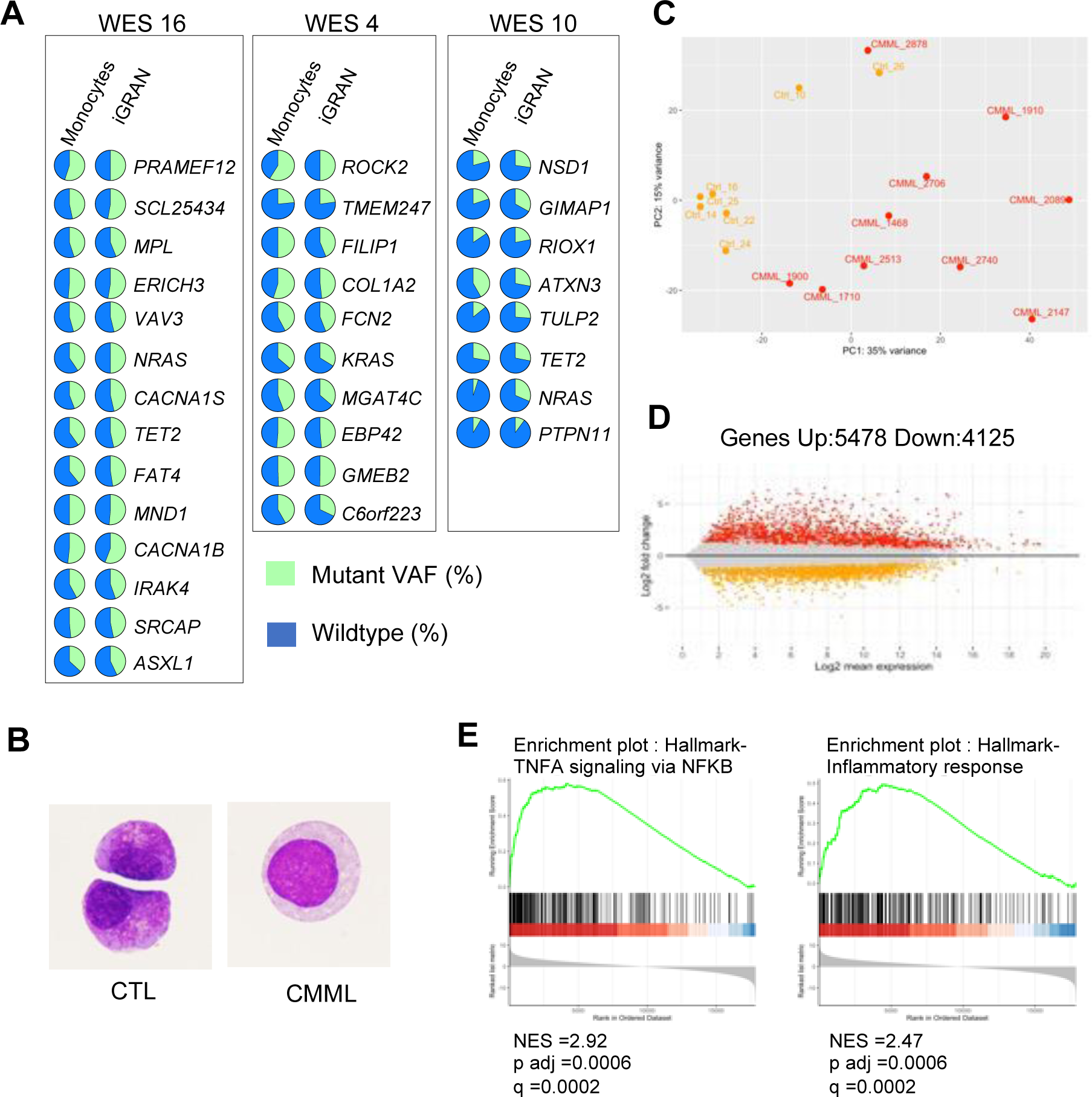
iGRANs are clonal, pro-inflammatory granulocytes. **A**. Variant allele frequency (VAF) of indicated somatic variants detected by whole exome sequencing (WES) of sorted monocytes (left) and iGRANs (right) in three CMML patients (16, 4, 10). **B**. May-Grünwald Giemsa staining of iGRAN sorted from a control donor (CTL) and a CMML patient sample, Magnification ×1000. **C-E**. Bulk RNA sequencing of sorted iGRANs collected from 7 healthy donors (CTRL) and 10 CMML patients; PCA of regularized logarithm (rlog) transformed data based on the top 500 varying genes using plotPCA function of DESeq2 package (C). MA-plot of differentially expressed genes between iGRANs collected from healthy donors and CMML patients (D). Gene set enrichment analysis (GSEA) of indicated hallmark pathways enriched in CMML vs CTL (E), Adjusted p and q -values are indicated on the graphs.

The gene expression profile of human G-MDSC distinguishes them from mature neutrophils or monocytes (28). This profile was recapitulated in clusters 5 and 8 that exhibited high expression of cell cycle genes such as *MKI67,* as well as high expression of *ARG1, MPO, S100A8, ANXA1, CYBB, and S100A12* genes and did not express the *TNF* gene (**Figure 3B**). This gene signature was validated by RT-qPCR analysis in sorted CD16^+^ neutrophils, CD14^+^ monocytes and CD15^+^CD16^-^ iGRANs collected from 8 CMML patients. iGRANs expressed significantly higher levels of *ARG1, S100A12* and *MPO*; similar levels of *S100A8*, *S100A9* and *MMP9*; and lower levels of *TNF* compared to mature neutrophils or monocytes (**Figure 3F**). These features are consistent with the hypothesis of G-MDSC (28,29).

The main characteristic of G-MDSCs is their ability to suppress immune cells. Consistent with the spectral flow cytometry analyses showing a significant inverse correlation between the iGRAN and CD4^+^ T-cell fractions (r=-0.49, p=0.01, **Supp Figure 4D**), conventional flow cytometry analysis showed an inverse correlation between the iGRAN and lymphoid cell fractions (obtained by WBC count). We also cultured PBMCs collected from untreated iGRAN-low (<14%) and iGRAN-high (≥14%) CMML patients for 4 days with anti-CD3 and anti-CD28 antibodies for T cell activation. iGRAN depletion from iGRAN-high, but not iGRAN-low PBMCs increased CD4^+^ or CD8^+^ T-cell proliferation. When a fixed ratio of sorted iGRANs was added to iGRAN depleted-PBMC, T-cell proliferation was strongly reduced in all situations, whatever the initial fraction of iGRANs in PBMCs (**Figure 3G, Supp Figure 4E-F**). Taken together, these results demonstrate that iGRANs are G-MDSCs.

### iGRANs are clonal cells with high inflammatory activity

G-MDSC were shown to be part of the leukemic clone in acute myeloid leukemia (30) and in chronic myeloid leukemia (31). Ambiguity remains in myelodysplastic neoplasms (32), and the question has never been addressed in CMML. The early clonal dominance depicted in this disease (8) left little room to the hypothesis of non-mutated granulocytic differentiation, but a subclonal expansion was possible. To explore the link between iGRANs and the somatic genetic abnormalities that characterize CMML, we performed whole exome sequencing of sorted iGRANs, monocytes and T cells from 14 untreated CMML patients. For each patient analyzed, the somatic variants identified in iGRANs matched those detected in monocytes with similar variant allele frequencies (VAFs) (**Figure 4A and Supp Table 6**). These results indicate that iGRANs belong to the leukemic clone and their accumulation is not related to subclonal coding region variants.

To characterize further CMML-associated clonal iGRANs, we compared them to sorted CD15^+^CD16^-^ cells collected from healthy donor by cytapheresis after mobilization. Cytological examination validated the immature morphology of collected cells, corresponding to promyelocytes or myelocytes, and revealed greater dysplasia in CMML iGRANs, which showed a loss of cytoplasmic granules and less condensed nuclear chromatin (**Figure 4B**). Transcriptomic profiles were studied in CD15^+^CD16^-^ cells sorted from 7 healthy donors and 10 untreated CMML patients. Gene expression in the 17 pooled samples was ranked, and genes such as *DEFA1, DEFA3, S100A9, MPO,* and *LYZ,* were identified as the most highly expressed genes of the transcriptomic profile (**Supp Figure 5A**), whether for CMML and control samples (**Supp Figure 5B**). Again, these analyses validated the immature and granulocytic phenotype of the sorted cells.

**Figure 5.**
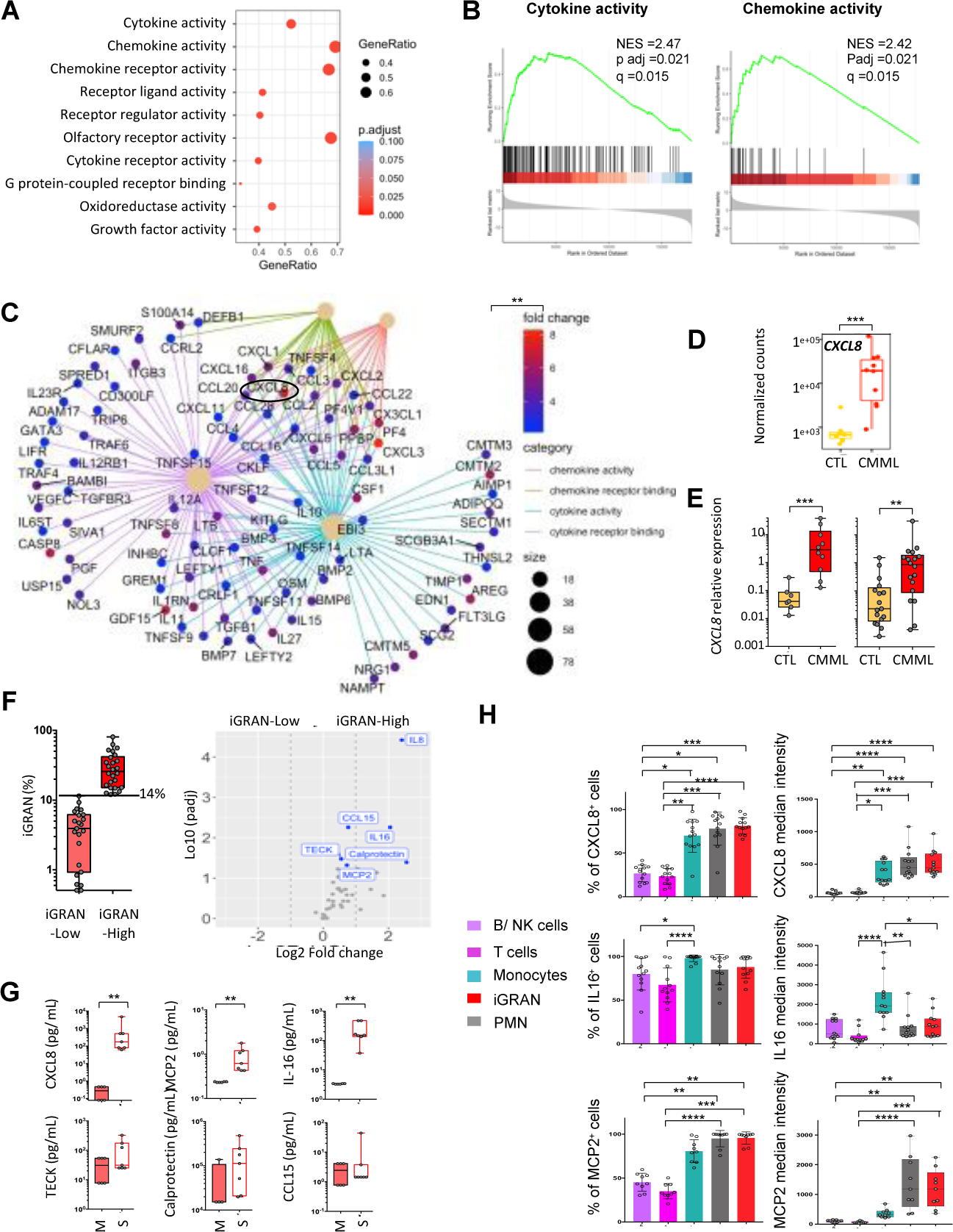
CXCL8 is one of the main cytokines released by CMML patient iGRANs. **A-D**. Bulk RNA sequencing of sorted iGRAN collected from healthy donors and CMML patients. Top10 nominal enrichment score of pathways involving up-regulated genes in CMML vs CTL cells according to GO molecular function (A). Gene set enrichment analysis (GSEA) of indicated hallmark pathways enriched in CMML vs CTL, adjusted p and q values are indicated on the graphs (B). Gene-concept network representation of enriched genes of 4 pathways (chemokine activity in pink, chemokine receptor binding in green, cytokine activity in blue, cytokine receptor binding in violet). Dot color, log2FoldChange. Size of beige central dots, number of core enriched genes (C). *CXCL8* mRNA expression (normalized counts) in CTL cells (N=7) and CMML iGRANs (N=10)(D). **E.** *CXCL8* mRNA expression assessed by RT-qPCR in healthy donor and CMML; left panels, samples used for RNA sequencing; right panel, independent cohort of iGRANs collected from 17 CTL and 18 CMML. Ct values normalized to *GAPDH*, *RPL32* and *GUS* genes. **F.** Volcano plot of cytokine and chemokine levels (N=44) measured in circulating plasma of 23 iGRAN-low compared to 26 iGRAN-high CMML (threshold ≥14%, iGRAN fraction in box plot for each group on the left panel). **G.** Indicated proteins were quantified in the supernatant of iGRANs (S), using culture medium (M) as a control. **H**. Intracellular cytokine production by B/NK cells, T cells, monocytes, iGRAN and neutrophils (PMN) is fresh peripheral blood samples collected from CMML patients. Left panels, fraction of cells expressing the studied cytokine; right panels, median fluorescence intensity for CXCL8 (N=13), IL16 (N=12) or MCP2 (N=9) in cells expressing the cytokine.

Principal component analysis (PCA) showed that patient and control cells mostly clustered separately (**Figure 4C, Supp Figure 5C**). Based on an adjusted P value of <0.05 and a log2-fold change >2, a total of 9,603 genes were found to be differentially expressed in CMML and control samples, of which 5,478 were upregulated (**Figure 4D**). Pathway analysis using gene set enrichment analysis (GSEA) for hallmarks identified 33 pathways significantly enriched in CMML iGRANs. The pathways with the highest enrichment scores were TNFA_signaling_via_NFKB, inflammatory response, and interferon response, whereas various metabolic pathways were downregulated (**Figure 4E, Supp Figure 5D**). These results indicate a proinflammatory status of clonal iGRANs in CMML patients.

### CXCL8 is the main cytokine secreted by iGRANs

GSEA molecular function analysis of RNA sequencing data showed the significant enrichment of gene sets associated with receptor binding and cytokine and chemokine activity in CMML-associated iGRANs (**Figure 5A, B**). Analysis of individual cytokine-encoding genes identified the *CXCL8* gene (*C-X-C motif chemokine ligand 8*, also known as *interleukin-8* or *IL-8*) as the most highly overexpressed (**Figure 5C, D**). *CXCL8* overexpression was confirmed by RT-qPCR analysis of two different sets of control and CMML cells (**Figure 5E**). Then, we measured the plasma levels of 44 cytokines and chemokines in iGRAN-low (N=23) and iGRAN-high (N=26) CMML patients and found that CXCL8 was the cytokine present at the highest level in the plasma of iGRAN-high CMML patients (**Figure 5F**). CCL15 (C-C motif chemokine ligand 15), IL-16, the S100A8/S100A9 heterodimer (also known as calprotectin), MCP2 (monocyte chemoattractant protein 2, also known as CCL8), and TECK (CCL25) were also detected at significantly higher levels in the plasma of the iGRAN-high patient group compared to the iGRAN-low patient group (**Supp Figure 5E**). Importantly, CXCL8, CCL15, calprotectin and IL-16 plasma levels correlated with the iGRAN fraction detected by flow cytometry in CMML peripheral blood (**Supp Figure 5F**). Of these cytokines, CXCL8, IL-16 and MCP2 were those detected at high concentrations in the culture supernatant of sorted CMML-associated iGRANs (**Figure 5G**). By performing intracellular staining of fresh peripheral blood samples including neutrophils, we detected these three cytokines in both mature (neutrophils) and immature (iGRAN) granulocytes. IL-16 was more abundant in monocytes that also produce CXCL8 but low levels of MCP2 (**Figure 5H**). A significant correlation was observed between the fraction of iGRAN in CD11b^+^CD33^+^ cells and CXCL8 median fluorescence intensity in mature neutrophils (**Supp Figure 5G**). Altogether, the ability of cells of the neutrophil lineage to produce CXCL8 may reflect the level of dysplasia in this lineage.

### CXCL8 inhibition restores the growth of wildtype CD34^+^ cells

In CMML, the majority of hematopoietic stem and progenitor cells (HSPC) are clonal, mutated cells, with a low number of residual wildtype cells. We evaluated the impact of the three main cytokines produced by iGRANs on healthy donor (cord blood and adult bone marrow) and CMML (bone marrow) CD34^+^ cells. In liquid culture, we observed a dose-dependent inhibition of cord blood CD34^+^ cell growth in the presence of CXCL8 (**Figure 6A**); a similar inhibitory effect was observed using adult healthy donor bone marrow CD34^+^ cells cultured with CXCL8 (**Supp Figure 6A**). In contrast, neither IL-16 nor MCP2 modified healthy donor CD34^+^ cell growth (**Supp Figure 6B**). Importantly, the three cytokines including CXCL8 failed to inhibit CMML CD34^+^ cell growth (**Figure 6A** and **Supp Figure 6B**). Together, these results indicated a specific inhibitory effect of CXCL8 on the growth of wildtype cells. Consistent with these results, the growth rate (33), calculated from the number of cells generated after 3 days in culture, was specifically decreased in healthy CD34^+^ cells in the presence of CXCL8. CMML cells had a similar average growth rate as compared to healthy CD34^+^ cells, although with higher inter-sample heterogeneity, and their growth rate remained unchanged in the presence of CXCL8 (**Supp Figure 6C**).

**Figure 6.**
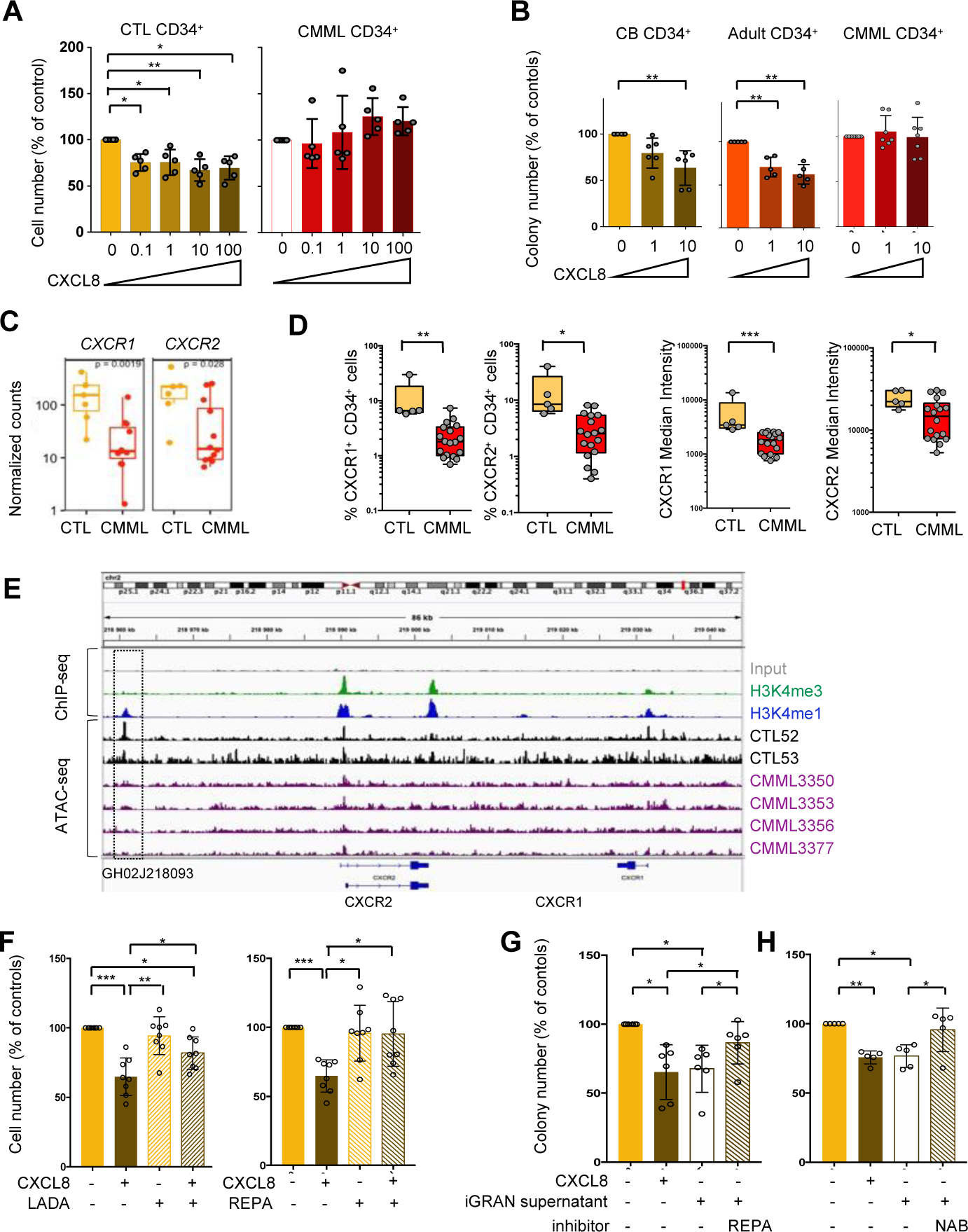
CXCL8 specifically inhibits the growth of wildtype CD34^+^ cells. **A.** Cell output of CMML CD34^+^ cells in liquid culture for 3 days in the presence of indicated doses of CXCL8 (ng/ml); ratio to untreated samples; mean ± SD, N=5 per group. **B.** Total colony output of CD34^+^ cells cultured in methylcellulose in the presence of indicated doses of CXCL8 (ng/ml) for 14 days. CB, cord blood (N=6); Adult, healthy donor bone marrow CD34^+^ cells (N=5) or CMML samples (N=7); ratio to untreated samples; mean ± SD. **C.** *CXCR1* and *CXCR2* mRNA expression assessed by RNA sequencing of CD34^+^ cells sorted from healthy donors (N=7) and CMML patient (N=12) bone marrow. **D.** Flow cytometry analysis of CXCR1 and CXCR2 at the surface of healthy donor (CTL, N=5) and CMML patient (N=18) CD34^+^ cells; left panels, fraction of cells expressing the studied receptors; right panels, within positive cells, mean fluorescence intensity of each receptor. **E.** *In silico* data for input (gray), H3K4me3 (in green) and H3K4me1 (in blue) ChIP-Seq obtained in human healthy CD34^+^ cells. Assay for Transposase-Accessible Chromatin (ATAC) sequencing experiments performed CD34^+^ cells obtained from two healthy donors (CTL, in black) and 4 CMML bone marrow samples (in purple). Rectangle: peak calling for CXCR2 enhancer. **F.** Cell output of healthy donor CD34^+^ in liquid culture in the absence or presence of CXCL8 10 ng/mL, or Ladarixin (LADA) 10 µM, or Reparixin (REPA) 10 µM. Mean ± SD, N=8. **G.** Total colony output of healthy donor CD34^+^ cultured in methylcellulose in the absence or presence of CXCL8 (10 ng/mL), or iGRAN supernatant, or reparixin (REPA, 10 µM); ratio related to untreated samples; N=6; mean ± SD. **H.** The same experiment was performed by using a CXCL8 neutralizing antibody (NAB, 5µg/mL); N=5 mean ± SD.

Methylcellulose colony formation assays also showed a decrease in colony number when healthy, cord blood or bone marrow CD34^+^ cells were seeded in the presence of CXCL8 whereas, again, the number of colonies generated by CMML CD34^+^ cells was not affected by the presence of CXCL8 (**Figure 6B**). This decrease affected CFU-GM (granulocyte-monocyte colony-forming unit) and CFU-E (erythrocyte-colony-forming unit) number, as observed by microscopic visualization (**Supp Figure 6D**) and validated by spectral flow cytometry analysis of the colony phenotype (**Supp Figure 6E**).

To explore further CXCL8-induced inhibition of wildtype CD34^+^ cell growth, we labeled these cells with cell trace to measure the number of cell division after 3 days in liquid cultures and calculate a proliferation index. CXCL8 did not modify the proliferation index of cord blood or bone marrow healthy CD34^+^ cells, suggesting that the difference in cell growth could be due to differences in cell death rate (**Supp Figure 6F**). In favor of this hypothesis, the decreased number of colonies in methylcellulose was associated with a trend for more Annexin-V^+^ cells, which was amplified by serial replating of the cells (**Supp Figure 6G**).

This lack of a response of CMML CD34^+^ cells to CXCL8 correlated with decreased expression of the CXCL8 receptors CXCR1 and CXCR2 at both the mRNA (**Figure 6C**) and protein (**Figure 6D**) levels. Using flow cytometry (gating strategy, **Supp Figure 6H),** we show that the fraction of CD34^+^ cells expressing CXCR1 and CXCR2 at the cell membrane and the median fluorescence intensity of these receptors are both significantly decreased in CMML compared to healthy donor samples. Searching for a mechanism that could account for the decreased expression of these receptors, we used *in silico* ChIP-Seq data performed in healthy CD34^+^ cells to pinpoint *CXCR1* and *CXCR2* transcription starting sites (TSS) using H3K4me3 marks and *CXCR2* gene enhancer (GH02J218093) using H3k4me1 marks. We then performed ATAC-seq experiments in CD34^+^ cells from healthy donors and CMML patients. ATAC-seq data showed a peak at *CXCR1* and *CXCR2* TSS in healthy donor CD34^+^ cells (in black). These peaks were hardly distinguished in CMML samples. Similarly, the peak identified at the *CXCR2* enhancer of healthy CD34^+^ cells was poorly detected in patient samples (**Figure 6E**). Together, these results suggested an epigenetic down-regulation of *CXCR1* and *CXCR2* in CMML CD34^+^ cells.

In an attempt to counteract the negative effect of iGRAN-derived CXCL8 on wildtype CD34^+^ cell growth, we checked the effect of pharmacologic inhibition of CXCL8 receptors, CXCR1 and CXCR2. Among the various small molecule CXCR1/2 antagonists that are being developed clinically, two were tested, namely ladarixin (4-[(2R)-1-oxo-1-(methanesulfonamide)])(34) and reparixin (*R*(–)-2-(4-isobutylphenyl)propionyl methanesulfonamide)(35). Consistent with our previous results, the presence of exogenous CXCL8 in culture decreased colony forming capacity of wildtype CD34^+^ cells (**Supp Figure 6I**) as well as their proliferation in liquid culture (**Figure 6F**), and the addition of ladarixin or reparixin in combination with CXCL8 restored the proliferation and the colony forming capacity of wildtype CD34^+^ cells, validating the efficacy of these drugs in this setting. iGRAN supernatant added to cell cultures had the same inhibitory effect as CXCL8 alone on CD34^+^ colony forming capacity, and blockade of CXCL8 using a CXCR1/2 inhibitor or a CXCL8 neutralizing antibody prevented the inhibitory effects of iGRAN supernatant (**Figure 6G-H**). Together, CXCL8 secreted by iGRANs that accumulate in the peripheral blood of CMML patients inhibits the growth of wildtype CD34^+^ cells while sparing CMML CD34^+^ cells in which CXCL8 receptors are down-regulated. These data suggest that, in the context of CMML, CXCL8 neutralization or CXCL8 receptor pharmacological inhibition might contribute to restoring the growth of residual wildtype CD34^+^ cells.

## Discussion

The overarching aim of this study was to refine the detection of immature granulocytes named iGRANs in the peripheral blood of CMML patients and to explore their pathogenic role in disease progression. We show that flow cytometry quantification of iGRANs among circulating myeloid cells provides an independent prognostic information. These cells behave as myeloid-derived suppressive cells and secrete large amounts of CXCL8, an inflammatory cytokine that specifically inhibits wildtype CD34^+^ cells. In contrast, CXCL8 does not affect leukemic CD34^+^ cells in which CXCL8 receptors are epigenetically downregulated. The ability of reparixin, an inhibitor of CXCL8 receptors, to counteract the negative effect of iGRAN supernatant on wildtype hematopoiesis suggests a strategy to slow down CMML evolution by re-expanding residual healthy hematopoiesis.

Flow cytometry analysis of peripheral blood cells, which supports CMML diagnosis by identifying an abnormal partition of monocyte subsets (2,21,22), is shown here to also be a powerful stratification tool by quantifying the iGRAN fraction or their absolute number. The cytological detection of immature granulocytes was suspected to define a poor prognostic subgroup of CMML patients (36), but the limited accuracy of IMC quantification on routine blood tests precluded its incorporation in the most used prognostic scores, which feature diverse combinations of age, cytopenias, and cytogenetic and molecular markers (24–26). A more precise and reproducible measurement of iGRAN fraction by flow cytometry identification of CD45^low^, CD11b^+^, CD14^-^, CD15^+^, CD16^-^, CD24^+^, CD33^+^, CD66b^+^, HLA-DR^-^ cells in the peripheral blood, with cutoff values at 14% of CD11b^+^CD33^+^ cells or 0.4×10^9^/L, may refine existing stratification scores. Besides, the accumulation of myeloid cells with a similar phenotype is a well-identified negative prognostic factor in multiple other pathological situations, including infection, trauma and cancer (37).

The accumulation of iGRANs in the peripheral blood of a fraction of CMML patients questions the mechanisms involved in their generation. The accumulation of iGRANs does not reflect subclonal genetic evolution as these clonal cells express all the genetic variants identified in monocytes without detecting additional genomic event. The recently identified role of the chromatin regulator Additional sex combs-like 1 (ASXL1) in neutrophil development, based on the neutrophilic dysplasia observed in an *Asxl1*-truncated zebrafish model (38) and the altered transcription program depicted in *Asxl1*-mutated mouse granulocyte progenitors (39) may account for the correlation between iGRAN excess and *ASXL1* gene mutation in CMML patients, both events being associated with a poor outcome (4,19,24–27). Some other disease features may contribute to the immunosuppressive phenotype of iGRANs. For example, the proliferative CMML subtype involves myeloid progenitor hypersensitivity to GM-CSF (9), a cytokine that promotes the generation of G-MDSCs in various other settings (37). Mutations in splicing regulator genes, that also correlate with IGRAN excess, could promotes the immunosuppressive activity of G-MDSCs by activating the NF-κB signaling pathway (40,41). Finally, iGRAN excess associates with anemia and lymphocytopenia, which may be related to the immunosuppressive potential of iGRANs (28) and their pro-inflammatory phenotype (42).

We show here that iGRANs are part of a dialog between clonal mature cells and HSPCs. When the occurrence of a somatic mutation in a single HSC leads to clonal outgrowth, mature myeloid cells from the clone commonly demonstrate an inflammatory phenotype and promote multiple diseases (43–45), including myeloid malignancies (46). In the context of CMML, monocytes have been shown to secrete cytokine-like 1 (CYTL1), which reduces monocyte apoptosis through an autocrine or paracrine pathway involving MCL-1 and the MAPK pathway (47), while MIF, which is released in the context of *TET2* truncation mutations, promotes the monocytic differentiation of HSPCs in a feed-forward loop (10). Here we show

an increased production of CXCL8 by the neutrophil lineage, correlated to the amount of iGRAN accumulation. The observation that the inflammatory cytokine CXCL8 specifically impacts wildtype CD34^+^ cell expansion and not CMML CD34^+^ is reminiscent of a zebrafish model of clonal hematopoiesis in which myeloid cells derived from mutant HSPCs secrete inflammatory cytokines that repress the growth of wildtype HSCs but do not impact mutated HSCs (48).

A specific decrease in the growth rate of wildtype CD34^+^ cells may increase the relative fitness of clonal CD34^+^ cells. The lack of *cxcl8* gene in mouse genome precludes the use of genetically modified mouse models to explore this hypothesis, while the clonal architecture of the disease with few residual wildtype cells and a growth advantage to mutated cells when undergoing differentiation (8) precludes the use of xenografted animals (47). As a heuristic tool to explore the significance of this effect, we used a mathematical model (49,50) of the HSC compartment to estimate their fixation time (i.e., the time for one mutant to take over the compartment) either with or without CXCL8. First, we noted that, without CXCL8, the relative fitness of CMML-mutated cells (calculated from growth rates) compared to healthy CD34^+^ cells *in vitro* is around 1, indicating very small or even no fitness advantage and resulting in very long fixation times. Second, for the 10 out of 19 patient samples in which CMML CD34^+^ cells had a higher fitness *in vitro* than healthy cells (fitness>1, **Supp Table 7**), the model predicts a drastic reduction in fixation time in the presence of CXCL8 (**Supp Figure 6J**). Last, the relative fitness of CMML CD34^+^ compared to healthy cells was always below 2 (range 1.08-1.67, **Supp Table 7**), and it is in this range that, independent of model parameterization, fixation times are most sensitive to changes in relative fitness (**Supp Figure 6K**). This suggests that, over the course of CMML development, the effect of CXCL8 on wildtype cells may speed up CMML clonal expansion by years to decades, which makes its inhibition an interesting clinical opportunity for patients with CMML.

Previous studies had detected high levels of multiple cytokines, including TNFα, IL-1β, IL-6, and CXCL8, in the circulating plasma and bone marrow supernatant of CMML patients, leading to heterogeneous patterns of inflammatory protein levels (12). To date, these patterns have failed to predict the clinical response to therapeutic agents such as ruxolitinib (51). Here, six cytokines were found to be overproduced in the peripheral blood of patients in the iGRAN-high CMML group, of which CXCL8 was at the highest level, showed a direct correlation with iGRAN quantification, and was detected in iGRAN supernatant and by intracellular staining. These results suggest that in the heterogeneous population of CMML patients, combining iGRAN quantification by flow cytometry with circulating CXCL8 level could define a subgroup of patients most likely to benefit from a therapeutic strategy targeting the CXCL8-mediated pathway.

In some diseases, mutant HSCs were shown to resist the chronic inflammation that otherwise triggers the exhaustion of nonmutated HSCs by switching from canonical to noncanonical NF-κB signaling (52). In the context of acute myeloid leukemia for example, IL-1 secreted by monocytes and myeloid blast cells promotes the growth and clonogenic potential of pathogenic CD34^+^ cells while suppressing colony formation by wildtype CD34^+^ cells (53). Another example is the ability of *JAK2*-mutated (54) and *TET2*-mutated (55) HSPCs to resist the suppressive effect of TNF on wildtype HSPCs. In CMML, the absence of impact of CXCL8 on clonal cells correlates with the downregulated expression of its receptors through a closed conformation of chromatin region at *CXCR1* and *CXCR2* loci, which enforces the role of epigenetic modifications in the phenotype of this disease (56).

Together, iGRAN excess quantified by flow cytometry appears to be a novel and independent prognostic factor that could improve the performance of existing stratification scores in CMML. Our results indicate that CXCL8 secreted by dysplastic granulocytes specifically inhibits wildtype CD34^+^ cells, which may give a competitive advantage to CMML-mutated cells that have lost CXCL8 receptor expression. By relieving CXCL8 selective pressure on wildtype HSPCs, reparixin, an orally available inhibitor of CXCR1 and CXCR2 could modulate clonal evolution and slow down the progression of the disease. If this effect of CXCR1/2 inhibitors is validated clinically, which may be tested in a close future, its activity could be secondarily enforced by combining such an inhibitor with hypomethylating agents that reduce cell dysplasia or with cell signaling targeting drugs that decrease cell proliferation.

## Methods

### Healthy donor and patient samples

Peripheral blood (PB) samples were collected after written informed consent with institutional review board approbations and before any treatment from patients with a diagnosis of CMML according to the WHO classification. PB films from 580 CMML collected from Mayo clinic (Rochester, MN, USA) were evaluated for immature myeloid cells (IMC [myelocytes, metamyelocytes, promyelocytes] ≥1%) (19). CMML samples of the learning cohort were collected between March 2015 and April 2019 from eight French centers and those of the validation cohort were collected independently between March 2015 and December 2021 from 10 French Centers (**Supp Table 4**). Control samples were routine tube remnants (age ≥65-years, Henri Mondor Hospital Créteil, France), remnants from cytapheresis in stem cell donors (Gustave Roussy, Villejuif, France) and buffy coats from blood donors (Age <65-years, Etablissement Français du Sang, Rungis, France). Bone marrow CD34^+^ cells from adult healthy donors were obtained from Lonza laboratories and the bone bank of Cochin hospital, and umbilical cord blood samples were collected from Saint-Louis hospital (AC-2016-2759, Paris).

### Cell sorting

PB samples collected on EDTA were processed within 24 hours. When indicated, Blood Cell Stabilizer (Cytodelics, Cytodelics AB) was mixed at a 1:1 ratio to 1mL of whole blood and transferred to −80°C freezer. In other cases, samples were centrifuged at 300 g for 5 min at room temperature (RT), plasma was collected, then peripheral blood mononuclear cells (PBMC) were isolated using Pancoll density centrifugation (Pan-Biotech, Dutscher). CD16^+^ neutrophils were sorted from the white cell layer directly above the red blood cells using immuno-magnetic microbeads (AutoMacs system, Miltenyi Biotech). PBMC were used for conventional flow cytometry or immuno-magnetic sorting of CD3^+^ T-cells, CD14^+^ monocytes or iGRANs (Classical Monocyte Cocktail, Miltenyi Biotec). Sorted cells (purity ≥90%) were stored at −80°C as dry pellets. iGRANs were centrifuged on microscope slides, dried for 1 hour at RT, and stained with May-Grünwald-Giemsa. Patient CD14^+^ DNA were subjected to next generation sequencing (NGS) for a myeloid panel (3). CD34^+^ cells were sorted by AutoMacs system and frozen in FBS-DMSO 10%.

### Spectral and conventional flow cytometry

Cryopreserved PB samples were thawed at 37°C and fixed before red blood cell lysis, as recommended by the manufacturer. Cells were washed and incubated with antibodies for 1 hour at 4°C, washed in BSA 1%/ EDTA 0.5M in PBS and analyzed on CyTEK Aurora flow cytometer (Cytek Biosciences). FCS files were exported using FlowJO software. Marker expression values were transformed using the auto-logicle transformation function. Phenograph clustering was performed using 28 markers and a number of nearest neighbors of 30. UMAP was run with a nearest neighbor of 15 and a min_distance of 0.2. Conventional flow cytometry analysis was performed on 200 µL of whole blood cells using a lyse no wash protocol (Versalyse lysing solution, Beckman Coulter) or on 2 ×10^6^ PBMC labeled and analyzed using a Fortessa cytometer (BD Biosciences) and the Kaluza software (Beckman-Coulter). For intracellular staining, whole blood (100µL) was diluted in 400 µL of complete medium (RPMI 1640 (Gibco) and incubated 3h with GolgiPlug (BD Biosciences). Cells were then stained at 4°C with antibodies. After red cell lysis (1-step Fix/Lyse solution, Invitrogen), cells were permeabilized with the permeabilization buffer (Invitrogen) and stained with anti-CXCL8-PE-Cy7 (BioLegend), anti-IL-16-PE (BioLegend) or anti-MCP2-eFluor 660 (BD Biosciences). Cell death was identified by analysis of cells stained with annexin V-(AnV-FITC) and propidium iodide (PI) antibodies before flow cytometry analysis as recommended (BD Biosciences).

### 3’ Single-cell RNA sequencing

PBMC were loaded onto a Chromium Single Cell Chip (10X Genomics), and captured mRNAs were barcoded during cDNA synthesis using the Chromium Next GEM Single Cell 3′ GEM, Library & Gel Bead Kit v3.1 (10X Genomics). Libraries were sequenced on NovaSeq 6000 (Illumina). Raw BCL-files were demultiplexed using bcl2fastq (version 2.20.0.422 from Illumina) and read quality control performed using fastqc (version 0.11.9). Reads were pseudo-mapped to the Ensembl reference transcriptome v99 (homo sapiens GRCh38 build with kallisto, version 0.46.2). The index was made with kb-python (version 0.24.4) wrapper of kallisto. Barcode correction using the whitelist provided by the manufacturer and gene-based reads quantification was performed with BUStools (version 0.40.0). Empty droplets were detected using the emptyDrops function from the dropletUtils package (version 1.10.3), barcodes with p-value < 0.001 (Benjamini–Hochberg-corrected) were considered for analysis. The count matrix was filtered to exclude genes detected in less than five cells, cells with less than 1500 UMIs or less than 200 detected genes, as well as cells with mitochondrial transcripts proportion higher than 20%. Cell cycle scoring was performed using the CellcycleScoring function of Seurat package (version 4.0.0), and the cyclone function of Scran (version 1.18.5). Doublets were discarded using scDblFinder (version 1.4.0) and scds (version 1.6.0). We manually verified that cells identified as doublets did not correspond to cells in G2M phase. Datasets were integrated using the Harmony method, merged using Seurat (version 4.0.4), and the SCTransform normalization method was used to normalize, scale, select 3000 Highly Variable Genes and regress out bias factors. The reduced PCA spaces were used as input for the HarmonyMatrix function implemented in Harmony package (version 0.1.0) where the batch effect (orig.ident) was regressed. The shared space output by harmony was used for clustering. The optimal number of dimensions was evaluated by assessing a range of reduced Harmony spaces using 3 to 49 dimensions, with a step of 2. For space, Louvain clustering of cells was performed using a range of values for the resolution parameter from 0.1 to 1.2 with a step of 0.1. The optimal space was the combination of kept dimensions and clustering resolution resolving the best structure (clusters homogeneity and compacity) in a UMAP. Marker genes for Louvain clusters were identified through a «one versus others» differential analysis using the Wilcoxon test through the FindAllMarkers function from Seurat, considering only genes with a minimum log fold-change of 0.5 in at least 75% of cells from one of the groups compared, and FDR-adjusted p-values <0.05 (Benjaminin– Hochberg method). UMAP visualization was done using Cerebro (version 1.2.2).

### Cytokine level measurements

Sorted iGRANs were cultured for 24 hours at 1 million/mL in RPMI medium. Supernatants were collected and centrifuged at 1500rpm for 10 min and stored at −80°C. Medium without iGRAN was used as control (N=3). Plasma aliquots were centrifuged at 1500 rpm for 15 min at 4°C, diluted 1:4 and analyzed using Bio-Plex ProTM Human Chemokine Panel 40-plex Assay (Bio-rad). Acquisitions and analyses were performed on a Bio-Plex 200 system with Manager 6.1 Software (Bio-rad), respectively. Soluble Calprotectin (1:100) and S100A12 (1:2) were measured using R-plex Human Antibody Sets (Meso Scale Discovery), a MESO QuickPlex SQ120 reader and the MSD’s Discovery Workbench 4.0. Each sample assayed twice, average value taken as final result.

### Lymphocyte proliferation assay

Ten million of PBMC were stained with anti-CD15, -CD16, - CD66b, -CD45 and -CD14 antibodies before sorting CD45^+^CD15^+^CD16^-^CD66b^+^CD14^-^ cells (iGRAN) using an Influx cell sorter (BD Biosciences). Total PBMC and iGRAN-depleted PBMC were suspended in Cell trace Violet (5 µM in PBS 1X, Thermo Fisher Scientific) for 15 min at 37°C, then plated in 96-well round bottom plates (1 million /mL in complete RPMI medium). When indicated, iGRANs were added to iGRAN-depleted PBMC (1:10 ratio). In cultures, T-cells were activated in wells coated with anti-CD3 (eBiosciences, clone OKT3) and anti-CD28 (eBiosciences, clone CD28.2) antibodies in IL-2-containg medium (0.01 ug/ml, Peprotech) for 4 days. Cells were labeled with LIVE/DEAD^TM^ Fixable Blue Dead Cell Stain Kit (Thermo Fisher Scientific) and antibodies and analyzed using a Fortessa.

### Cell culture and reagents

CD34^+^ cells were cultured for 72 h at 0.75 × 10^5^ cells/mL in complete MEM-alpha medium, using Stem cell factor (SCF, 50 ng/mL), interleukin-3 (IL-3, 10 ng/mL), thrombopoietin (TPO, 10 ng/mL) and FMS-like tyrosine kinase 3 (FLT-3, 50 ng/mL mL) in the absence or presence of CXCL8 in a 37 °C incubator with 5% CO2. All cytokines were from Peprotech. MCP2 was from thermofisher, IL16 from BioTechne. Cells were counted after Trypan blue staining. For methylcellulose assays, CD34^+^ cells were plated in duplicate at 500 cells with 1mL complete methylcellulose (MethoCult™ H4034 Optimum, Stem cell) with indicated doses of CXCL8. Colonies were enumerated and phenotyped at day 14. When indicated, CXCR1/2 inhibitors Reparixin and Ladarixin (MedChemTronica and Clinisciences, respectively), dissolved in dimethylsulfoxide (DMSO), or CXCL8 neutralizing antibody (MAB208) and Mouse IgG1 isotype control (Bio-Techne) were used.

### RNA extraction, RT-qPCR analysis

Total RNA was obtained from frozen dry pellet of sorted CD14^+^, CD3^+^ using Trizol™ Reagent (Thermo Fischer Scientific) and Direct-zol™ RNA Miniprep (Zymo research). For iGRAN, total RNAs were extracted using RLT buffer (Quiagen) and Trizol™ LS Reagent. Precipitated RNA was purified on a “mini-RNA” column (RNeasy Mini Kit from Qiagen), quantified on Nanodrop (Spectrophotometer ND-1000) and stored at −80°C. Total RNA was reverse transcribed with SuperScript IV reverse transcriptase with random hexamers (Thermo Fisher Scientific). Real-time quantitative polymerase chain reaction (RT-qPCR) was performed with AmpliTaq Gold polymerase in an Applied Biosystems 7500 thermocycler (Thermo Fisher Scientific). Primer sequences are available upon request.

### Bulk RNA sequencing

RNA integrity (RNA Integrity Score ≥ 7.0) was checked on the Fragment Analyzer (Agilent) and quantity was determined using Qubit (Invitrogen). SureSelect Automated Strand Specific RNA Library Preparation Kit was used with the Bravo Platform. Briefly, 100 ng of total RNA sample was used for poly-A mRNA selection using oligo(dT) beads and subjected to thermal mRNA fragmentation before convertion into double stranded DNA for library preparation. Final libraries were bar-coded, purified, pooled and paired-end sequenced on Novaseq-6000 sequencer (Illumina) at Gustave Roussy. Raw reads were mapped to hg19 genome with Tophat2 (v2.0.14) / Bowtie2 (v2.1.0). The number of reads per gene (GENECODE gene annotation v24lift37) was counted using HTSeq (0.5.4p5) and DESeq2 (v1.10.1) package was used for differential gene expression analysis. Gene Set Enrichment Analysis was performed using enrichplot package with a number of Permutation: 10 000, min Gene Set Size: 20, max Gene Set Size: 800, pvalueCutoff 0.05.

### Whole Exome Sequencing (WES)

DNA collected from sorted CD14^+^, CD3^+^ and iGRANs was assayed on Nanodrop, and 200 ng genomic DNA were sheared with the Covaris E220 system (LGC Genomics / Kbioscience). Fragments were end-repaired, extended with an “A” base at the 3’ end, ligated with paired-end adapters with the Bravo platform (Agilent) and amplified for ten cyclesFinal libraries were paired-end sequenced (2 x 100 bp reads) using Illumina NovaSeq-6000 sequencer. Somatic variants were detected in monocytes and iGRANs, using CD3 T-cells as a control, and validated on IGV software.

### Statistical analysis

Participants’ characteristics were reported as numbers and percentages for categorical variables, mean and standard deviation (normal distribution) or median and interquartile range (skewed distribution) for continuous variables. Differences between groups were compared using Mann-Whitney test (2 groups) or Kruskal-Wallis test (>2 groups). Correlation between clinical and biological parameters were assessed with the Spearman method. The p-values of the tests are expressed as follows: * p<0.05, ** p<0.01, *** p<0.001, **** p<0.0001. In the absence of precision, the test is not significant. A Cox proportional hazards model was used to adjust the effects of iGRAN fraction (%) or absolute number (×10^9^/L) as a continuous variable on overall survival (OS), defined as the time between the date of diagnosis and the date of death, whatever the cause (for patients who remain alive, OS was censored on the date of last follow-up), and event-free survival (EFS), defined as the time between the date of diagnosis and date of AML transformation or death due to any cause whichever occurs first (for patients who remain alive without AML transformation, EFS was censored on the date of last follow-up. The variables included in the Group Francophone des Myélodysplasies score (GFM) (24) were used in the multivariate models (Age, WBC >15x 10^9^/L, hemoglobin <10g/L, platelets < 100 x 10^9^/L, *ASXL1* mutations). The optimal cut-points were computed using maximally selected log-rank statistic (maxstat R package)(57,58) for overall survival to define two prognostic groups, and the Kaplan-Meyer method was used for survival curves (comparisons with log-rank tests). SAS 9.4 (SAS Institute Inc. Cary, NC) and R version 4.0.5 (R Foundation for Statistical Computing) softwares were used.

### Mathematical modelling

Evolution of N cells in the stem cell compartment was modelled using a Moran process (49) in which, at every iteration, one cell divides and one to die (can be the same cell), thus keeping the population size constant. The probability for a cell *i* to divide is proportional to its fitness *f_i_*

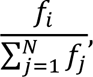

and the probability for a cell to die is

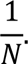

To model CMML, one malignant cell with relative fitness, r>1, is introduced at time *t=0* into a pool of N-1 identical healthy cells, each with fitness 1. For an advantageous mutated cell that expands to take over the whole cell compartment, the fixation time can be approximated using the following mathematical expression

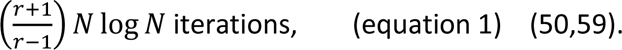

We assume that the whole compartment turns over in time 1/division rate, and hence the time for one iteration is 1/(division rate*N).

The model was parameterised using *in vitro* experimentally-derived values (for r) and values from the literature (for N and the division rate). In the basic scenario, we choose N=100,000 (60) and division rate 1/yr (61), Watson et al. (60) estimate Nτ≍100,000yrs where 1/τ is the self-renewal/differentiation rate, with lower and upper bounds on N of 25,000 and 1.3 million respectively, and τ<4yrs. We computed the growth rates of adult healthy bone marrow (N=3) and CMML (N=19) CD34^+^ cells by performing liquid cultures experiments in the absence or presence of CXCL8 (10ng/ mL) using an exponential model (33).

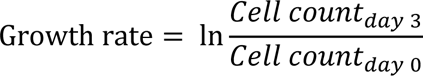

We calculated the fitness of every CMML sample as the ratio of the growth rate of each individual CMML sample to the mean growth rate measured in healthy cells, in the absence and presence of CXCL8. The mean growth rate of the 3 healthy samples used to compute CMML r was 1.17 +/- 0.13 (range 1.07-1.32) with CXCL8 and 1.54 +/- 0.09 (range 1.44-1.59) without CXCL8. As the mathematical expression for fixation time (equation 1) requires r>1, it was applied to compute the fixation time in CMML samples for which the fitness of CD34^+^ cells was higher than that measured in healthy samples (10 samples out of 19, r range 1.07-1.67).

## Author contributions

PD, MW, AG, MM, AI, VM acquired data. AP performed the statistical analysis. AML performed the mathematical calculations. AF, PR performed spectral flow cytometry analysis. RC, AR, MP analyzed data. VL, BB, OWB, CB, TB, CW, RI, PF, ST, GE provided samples. FP, EEM, ND, LP conducted experiments and analyzed data. LL analyzed bulk RNAseq experiments, conducted experiments for mathematical model and analyzed data. DSB and ES designed research studies, conducted experiments, analyzed data and wrote the manuscript.

## Acknowledgments

We gratefully acknowledge W. Vainchenker for helpful scientific discussions, J-E. Martin, A. Arbab, A. Dubuisson and B. Marteyn for their technical support and appreciate the constant support of the Groupe Francophone des Myelodysplasies (GFM). We thank J Lafosse for collecting clinical annotations, CYBIO platform and Gustave Roussy Bioinformatics and Cytometry platforms.

## Financial supports

The team is labelled by the Ligue contre le Cancer (to FP). We received grants from Institut National du Cancer (PRT-K Myelomono 2 to ES), Association Laurette Fugain, Cancéropole Ile-de-France, and SIRIC SOCRATE (to DSB), and ‘Taxe d’ apprentissage’ program (TA2019 to C.W. and TA2021 to V. M.).

## Notes

### Competing Interest Statement

The authors have declared no competing interest.

