## Supplemental material for "CXCL8 secreted by immature granulocytes inhibits wildtype hematopoiesis in chronic myelomonocytic leukemia"

### Deschamps et al. Supplemental Figure 1

**Supplemental Figure 1. Spectral flow analysis of peripheral blood cells. A.** Left, overall survival of 580 CMML patients using immature myeloid cells (IMC) cutoff (Yes, ≥ 1% of white blood cell, WBC). Middle, univariate analysis of IMC impact on overall survival; N=580 patients (IMC < 1%, N=229; IMC ≥ 1% WBC, N=351). Right, multivariate analysis built on 466 patients with available informations on IMC fraction, bone marrow blast cell count (CMML-2) and WBC count (MP-CMML. **B**. Cell surface marker expression in the non-supervised UMAP analysis of 37 samples (controls, N=10; CMML, N=27); **C-J.** Gating strategy for cell subset analysis in $CD45^+$ cell population. Debris and doublets were discarded by gating on “Cells” (**C**), “Singlets” and “Singlets bis” gates (not shown). Remaining red cells were excluded and $CD45^+$ cells were selected on a dot plot of CD45 vs SSC (**D**). $CD15^+$, $CD66b^+$ granulocytes were isolated in a “Neutro” gate (**E**) and separated into mature $CD15^+$, $CD16^+$ and immature $CD15^+$, $CD16^-$ neutrophils (not shown). Other hematopoietic cells in a “No Neutro” gate, $CD15^-$, $CD66b^-$ (**E**) include lymphoid populations, roughly isolated as expressing CD3 or CD19 or CD7 or NKG2D ($LIN^+$) using LIN vs HLA-DR dot plot, separating B-cells ($LIN^+$, $HLA\text{-}DR^+$) from NK / T cells ($LIN^+$, $HLA\text{-}DR^-$) (**F**). B-cells were further detected on a CXCR4 vs CD45RA dot plot (**G**), NK cells on a CD56 vs CD5 dot plot (**H**) and T-cells as $CD5^+$ cells, $CD4^+$ and $CD8^+$ T lymphocytes (**I**). With this exclusion gating strategy, remaining cells could be considered as monocytes and dendritic cells and monocyte subsets “Mono Pop” were separated from Dendritic cells “DC Pop” on a CD89 vs CD88 dot plot (**J**). **K.** Monocyte subsets in the “Mono pop” gate were separated on a CD14 vs CD16 dot plot into classical (cMO in blue), intermediate (iMO, in yellow) and non-classical monocytes (ncMO, in purple)(left panels) and the fraction of each subset was calculated in controls (CTL, N=10) and CMML (N=27) samples (right panels).

### Deschamps et al. Supplemental Figure 1

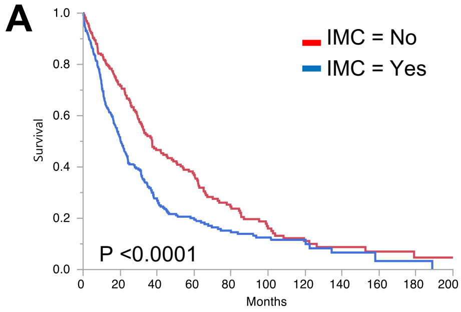

**Univariate analysis**

| Group | Median Time (Months) |
|---|---|
| IMC= No | 37.3 |
| IMC=Yes | 20.5 |
| Combined (n=580) | 26.2 |

**Multivariate analysis**

*Hazard Ratios (HR) and 95% Confidence Intervals*

| Analysis | Hazard ratio | p.value | 95% CI |
|---|---|---|---|
| IMC (yes) | 1.1563 | 0.2170 | 0.9182-1.4561 |
| Blast classification (CMML-2) | 1.0841 | 0.6716 | 0.7462-1.5751 |
| Proliferative form (MP-CMML) | 1.6546 | < .001 | 1.2974-2.1102 |

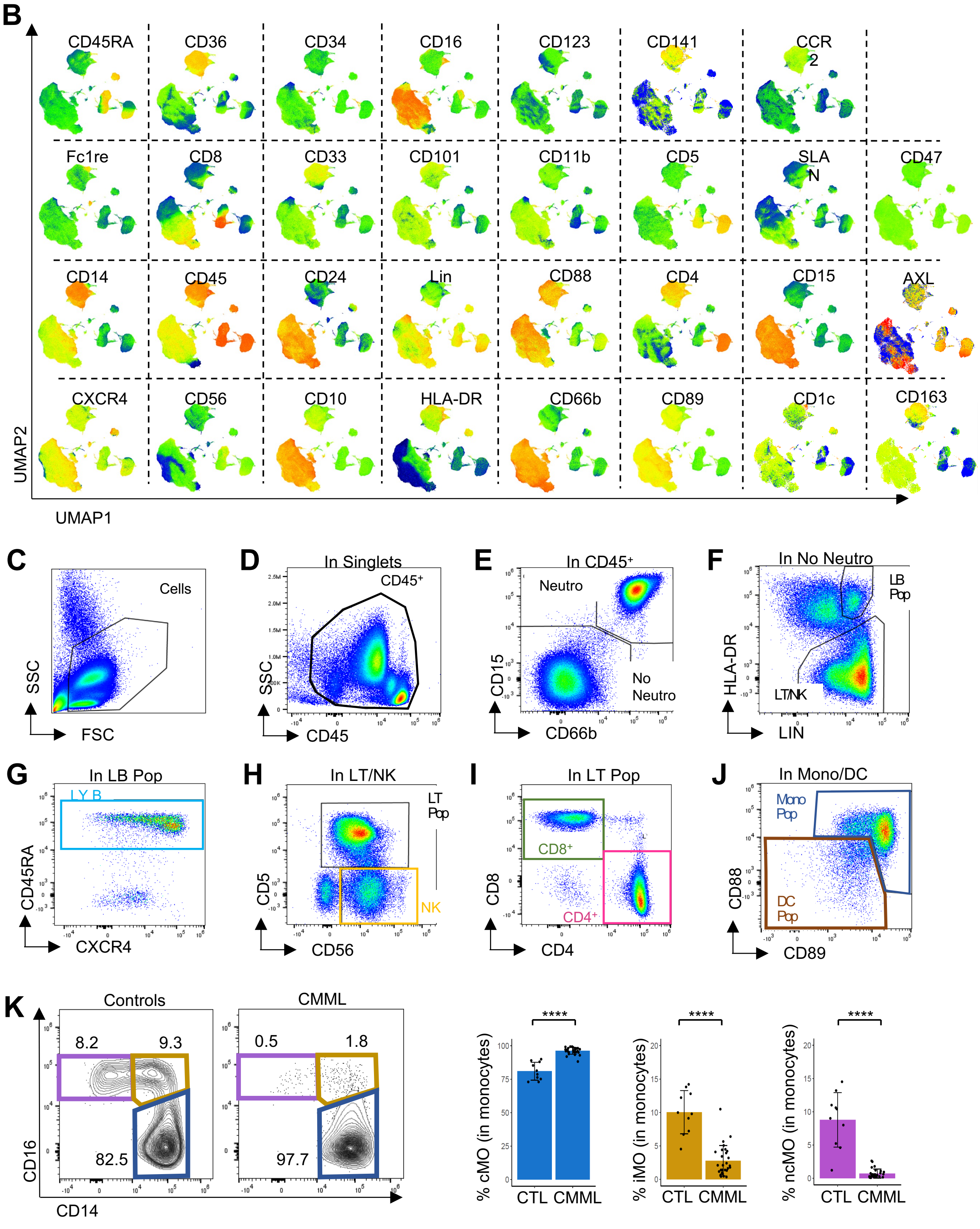

### Deschamps et al. Supplemental Figure 2

**Supplemental Figure 2. Conventional flow detection of iGRAN in CMML peripheral blood.** **A.** Representative flow plots showing the presence of iGRANs in the peripheral blood before (whole blood) and after low-density separation. **B.** Ratio of PMN ($CD15^+$, $CD16^+$ cells, upper panel, in grey) or iGRAN ($CD15^+$, $CD16^-$, lower panel, in red) to $CD45^+$ cells before and after low density separation. N=8. Student paired test. **C-J.** Step by step gating strategy to quantify iGRAN fraction after low-density separation of peripheral blood cells; forward (FSC) and side scatter (SSC-A) are used to discard debris and dead cells (**C**) and doublets (**D**); $CD45^+$ cells were selected on a CD45/ SSC plot (**E**); lymphoid cells were selected with a pool of Lin antibodies (anti-CD3, CD-19 and CD-56 in the same fluorescence channel, $LIN^+$) (**F**); remaining mature neutrophils were selected on a CD16/ SSC plot (**G**); total myeloid cells were selected as $CD33^+$, $CD11b^+$ cells in the "$LIN^-$ / Mature $neutro^-$" population (**H**); among the $CD33^+$, $CD11b^+$ myeloid cells, monocytes were identified as $HLA\text{-}DR^+$ cells on a HLA-DR vs SSC plot (**I**); iGRANs were identified by $CD15^+$ expression in the $HLA\text{-}DR^-$ population (**J**). **K.** Spearman correlation between iGRAN fraction calculated in $CD11b^+$, $CD33^+$ and in $CD45^+$ populations. **L.** iGRAN fraction was serially measured, up to three times over a period of one year, in CMML patients (N= 17).

#### Deschamps et al. Supplemental Figure 2

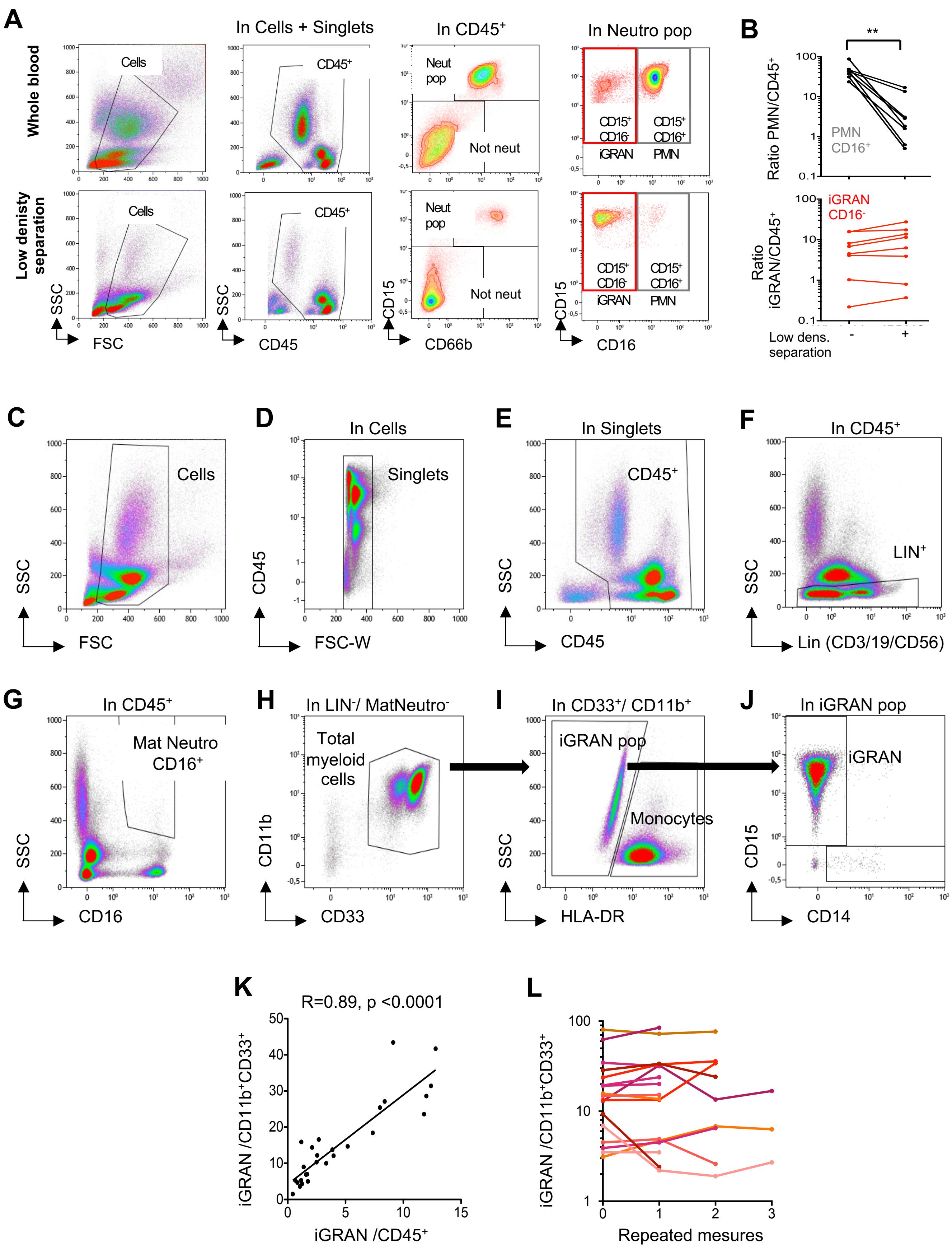

### Deschamps et al. Supplemental Figure 3

**Supplemental Figure 3. Relationship between iGRAN fraction, CMML features and patient outcome in the learning and the validation cohorts. A-G.** Learning cohort. **A.** Spearman correlation between iGRAN fraction and patient age. **B.** iGRAN fraction in female (F) and male (M) CMML patients. **C.** iGRAN fraction according to the presence or absence of cytogenetic abnormalities. **D.** Spearman correlation between iGRAN fraction and mature neutrophil fraction (PNN, left panel), monocyte fraction (middle panel) and platelet count (right panel). **E.** iGRAN fraction according to WHO 2016 CMML subtypes. **F.** iGRAN fraction according to the wildtype (WT) or mutated (Mut) status of indicated genes. **G.** Event-free survival (left) and overall survival (right) of CMML patients with high (≥0.4x$10^9$/L, N=63 in red) and low (<0.4x$10^9$/L, N=91 in blue) number of circulating iGRANs, log-rank test. **H-K.** Validation cohort. **H.** iGRAN fraction in $CD11b^+CD33^+$ population (left panel) and iGRAN absolute number (right panel) measured by flow cytometry in the peripheral blood of a validation cohort of CMML patients (N=160) compared to age-matched controls (N=64). **I.** iGRAN fraction in dysplastic (MD) and proliferative (MP) CMML subtypes. **J.** Spearman correlation between iGRAN fraction and immature myeloid cell (IMC) fraction, white blood cell count (WBC), hemoglobin level and lymphocyte fraction in the peripheral blood of CMML patients. **K.** Event-free survival and overall survival of CMML patients included in the validation cohort, using the cutoffs values calculated within the learning cohort. CMML patients with high (≥14%, N=34, in red) vs low (<14%, N=76, in blue) iGRAN fraction (left); CMML patients with high (≥0.4x$10^9$/L, N=27 in red) vs low (<0.4x$10^9$/L, N=79 in blue) number of circulating iGRANs (right), log-rank test.

### Deschamps et al. Supplemental Figure 3

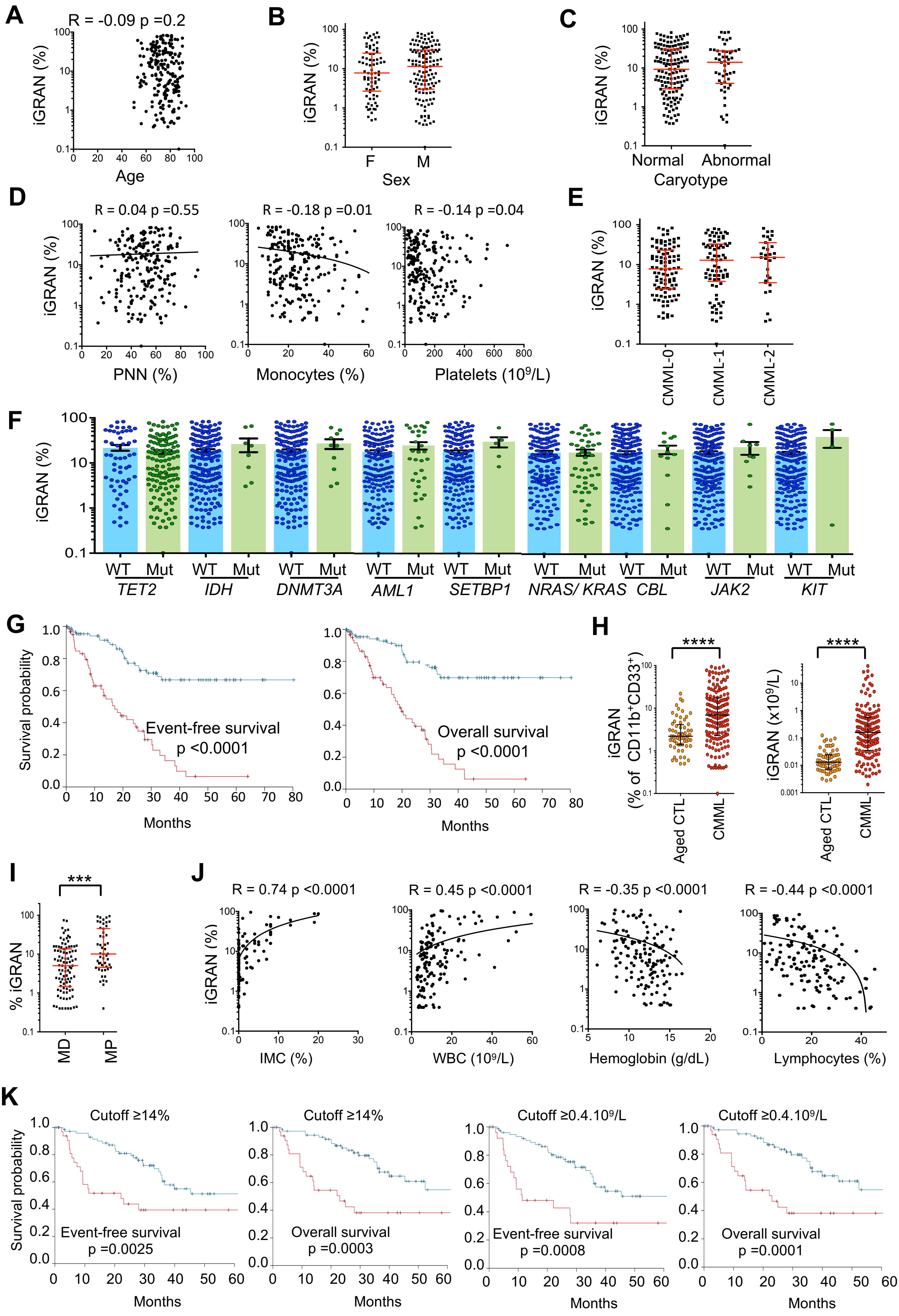

### Deschamps et al. Supplemental Figure 4

**Supplemental Figure 4. Molecular and functional characterization of iGRANs. A.** Heatmap displaying the scaled expression patterns of top marker genes within each cluster shown on Figure 3B. Scales indicate the intensity of expression. **B.** Expression level of indicated genes (low expression in yellow to high expression in red) in clusters identified on the UMAP shown on Figure 3B. **C.** Unsupervised clustering of cells in 3D (resolution 0.5 showing 15 main clusters). The dashed lined indicates ITGAM$^{+}$ myeloid population selected to calculate iGRAN fraction. **D.** Spearman correlation between iGRAN and CD4$^{+}$ T-cell fractions quantified by spectral flow cytometry. **E.** Proliferation assay performed in non-activated and anti-CD3/anti-CD28 activated T-cells; Forward (FSC) and side scatter (SSC-A) were used to discard debris. Dead cells and doublets were excluded using a live dead blue staining before selecting CD3$^{+}$ cells using FSC/ CD3 dot plot, then separating CD4$^{+}$ and CD8$^{+}$ T-cells. **F.** Examples of dot plots showing T cell proliferation measured by cell trace violet dilution in conditions described on Figure 3G.

### Deschamps et al. Supplemental Figure 4

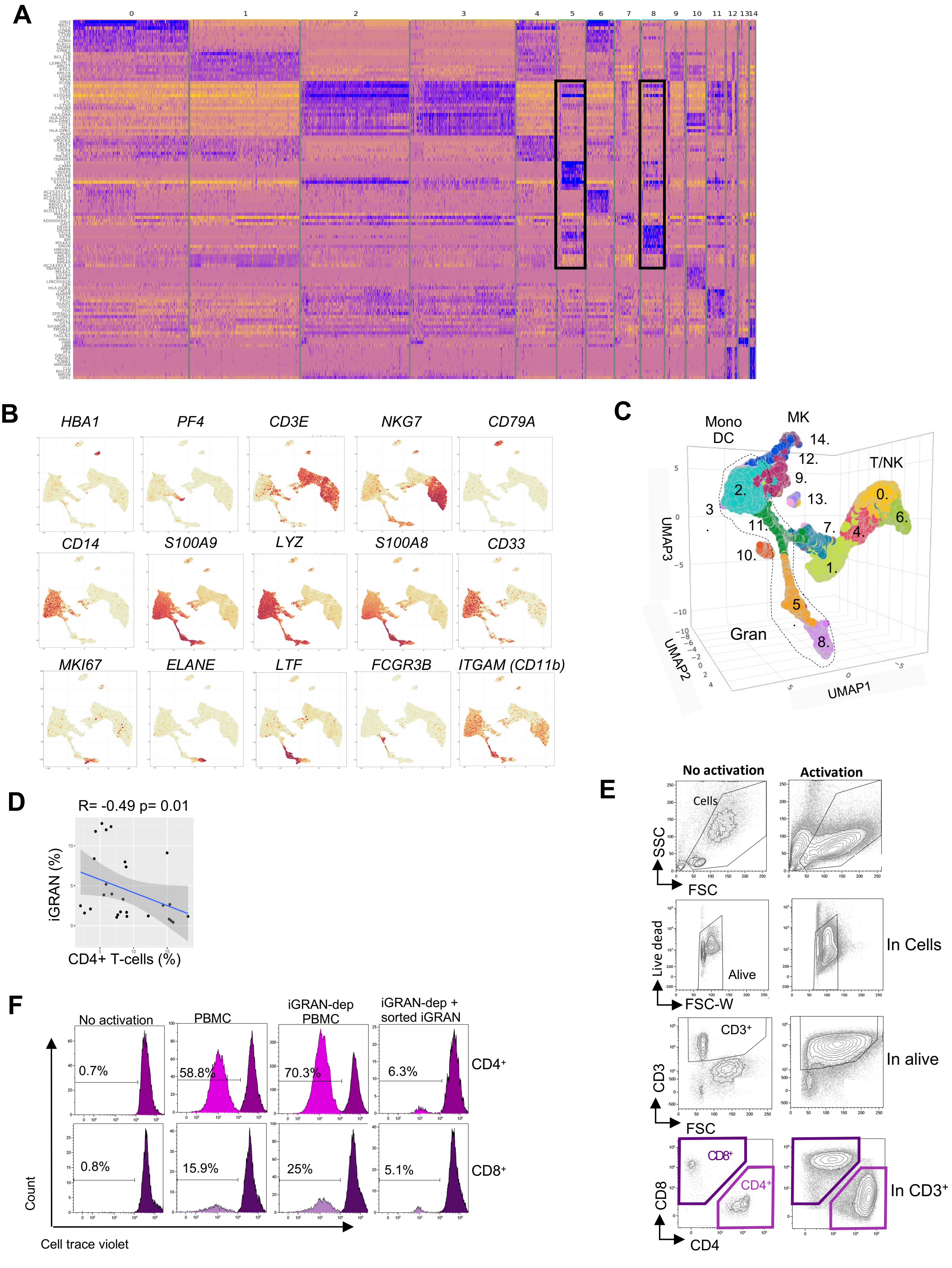

### Deschamps et al. Supplemental Figure 5

**Supplemental Figure 5. Sorted iGRAN RNA sequencing and cytokines associated to their accumulation in CMML patient peripheral blood.** iGRANs were sorted from healthy donor (cytapharesis, N=7) and CMML patient (N=10) peripheral blood. **A.** Mean expression of genes (using DESeq2 normalized counts) in the 17 samples sorted by decreasing order. Examples of genes highly expressed by iGRANs are in red and those highly expressed by monocytes in blue. **B.** Expression of genes (normalized counts) typically identified in iGRANs collected from controls (CTL) and CMML patients. **C.** Heatmap of sample-to-sample distances based on the rlog transformed counts. **D.** Ridgeplot representing gene set enrichment analysis (GSEA) of pathways distinguishing CMML from control iGRANs ($p<0.05$). **E.** Circulating levels of indicated cytokines in 23 iGRAN-Low and 26 iGRAN-High (cutoff ≥14%) CMML patients. **F.** Spearman correlation between iGRAN fraction in myeloid cells and plasma level of indicated cytokines. **G.** Spearman correlation between iGRAN fraction in myeloid cells and CXCL8 median fluorescence intensity in neutrophils (as shown in Figure 5H).

### Deschamps et al. Supplemental Figure 5

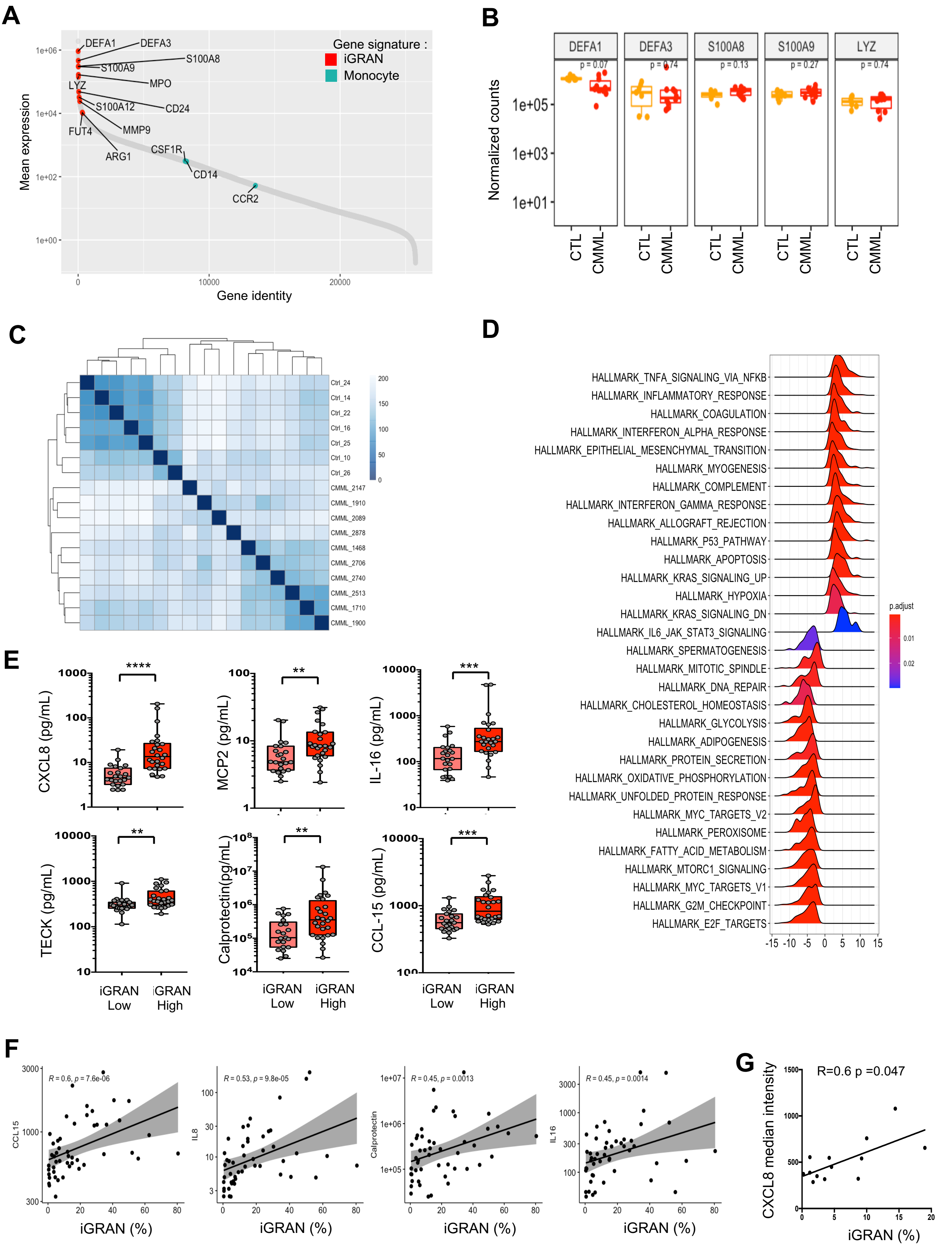

### Deschamps et al. Supplemental Figure 6

**Supplemental Figure 6. CXCL8 blockade restores CD34$^+$ wildtype growth. A.** Cell output of healthy donor bone marrow CD34$^+$ in liquid culture for 3 days in the presence of 10 ng/mL CXCL8; ratio to untreated samples; mean ± SD, N=6. **B.** Cell output of WT or CMML CD34$^+$ cells in liquid culture for 3 days in the absence or presence of 10 ng/mL CXCL8, IL-16 or MCP2; ratio to untreated samples; mean ± SD, N=6 per group**. C.** Growth rate (LN(number of cells at day 3 / number of cells at day 0)) of healthy donor bone marrow CD34$^+$ (N=3) or CMML CD34$^+$ cells (N=19) in liquid culture for 3 days in the absence or presence of 10 ng/mL CXCL8. **D.** Cord blood (CB, N=6 upper panels) and healthy donor bone marrow (adult, N=5 lower panels) CD34$^+$ were cultured in methylcellulose for 14 days to generate granulo-monocytic (CFU-GM, left panel) and erythroid (CFU-E, right panel) colonies in the absence or presence of indicated concentrations of CXCL8. Output of treated relative to untreated cells. Mean ± SD. **E.** Non-supervised UMAP of spectral flow cytometry analysis of cells generated in methylcellulose at day 14 in the absence or presence of CXCL8 (10 ng/mL, N=5 independent experiments). Left: UMAP analysis separating CD45$^-$, CD11b$^-$, CD36$^+$ erythroid cells from CD45$^+$, CD11b$^+$, CD14$^+$ monocytes and CD45$^+$, CD11b$^+$, CD15$^+$, CD66b$^+$ granulocytes; middle: projection of indicated markers on UMAP; right : fraction of erythroid or granulo-monocytic cells, ratio to untreated samples. **F.** Proliferation index measuring the number of divisions of CD34$^+$ cells collected from WT donors (cord blood, CB, N=3 or adult bone marrows, Adult, N=4), in presence or absence of CXCL8. **G.** Total colony output of healthy donor CD34$^+$ culture in methylcellulose in presence or absence of CXCL8 (10 ng/mL) during serial replatings (plating 1 or 2, left panel). Percentage of dead cells measured by flow cytometry (Annexin-V$^+$/ propidium iodide $^{-/+}$ ; right panel) (N=6). **H.** Gating strategy to explore CXCR1 and CXCR2 expression at the surface of CD34$^+$ cells. **I.** Total colony output of healthy donor CD34$^+$ cell culture in methylcellulose in presence or absence of CXCL8 (10 ng/mL) and a CXCR1/2 inhibitor [left panel, ladarixin (LADA, 10 µM); right panel, reparixin (REPA, 10 µM)]; ratio related to untreated samples; Mean ± SD; N=6. **J-K.** Fixation time (in years) estimated using a Moran birth-death process, assuming 100,000 HSCs and testing 6 different HSC division rates (from 0.5 to 3) **J.** Fixation time estimated from our growth rate data (panel C), for 10 CMML samples with a growth rate superior to the mean of CTRL growth rate, faceted by division rate, in presence (maroon) or absence (red) of CXCL8. **K.** Fixation time as a function of relative fitness, for relative fitnesses from 1.07 to 5 at steps of 0.01, faceted by division rate. Red points: the 10 CMML samples of panel J, in absence of CXCL8.

### Deschamps et al. Supplemental Figure 6

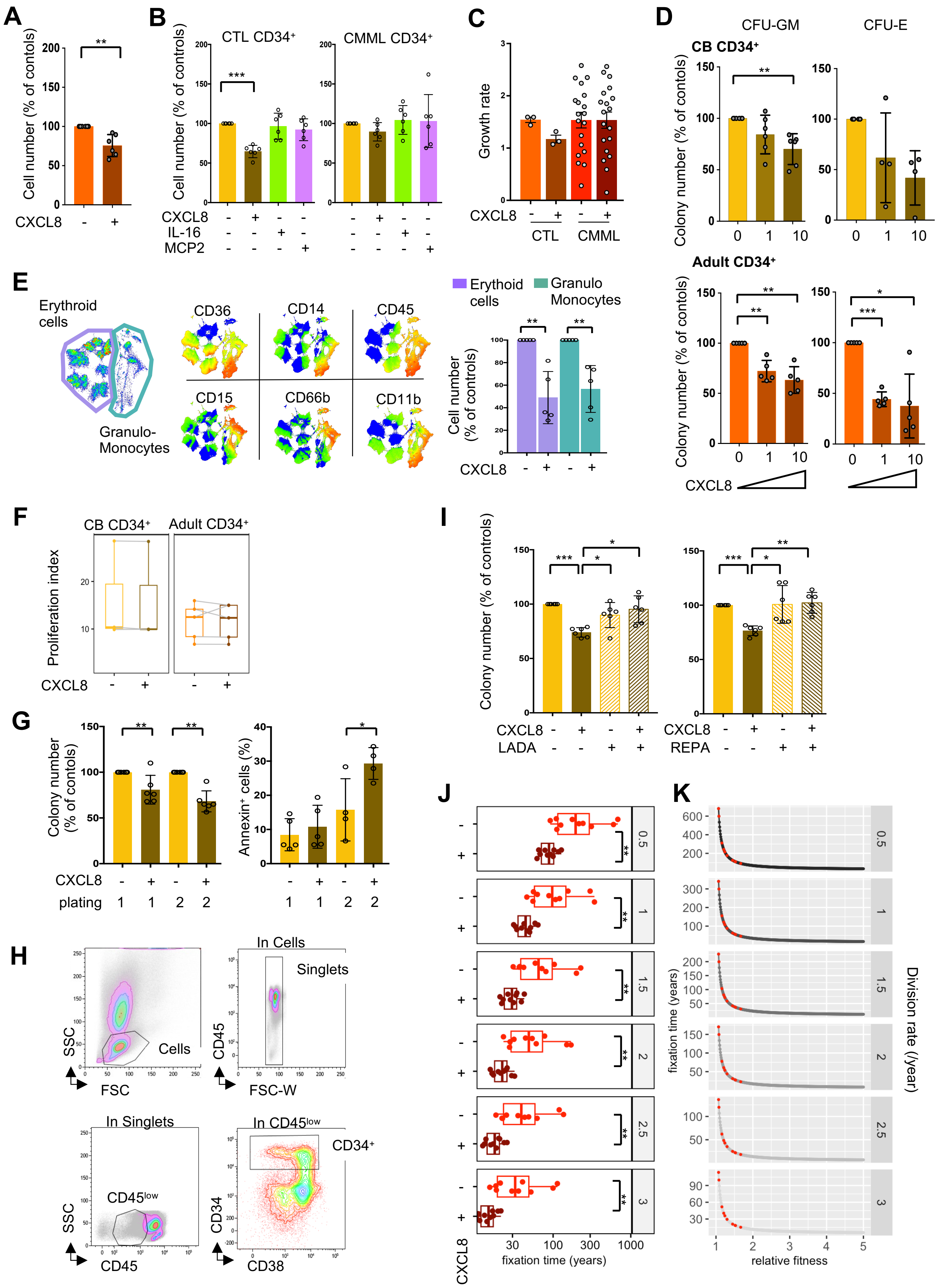

**Supplementary Table 1.** Clinical and laboratory characteristics of CMML patients stratified by the presence or absence of circulating immature myeloid cells (IMCs) in the peripheral blood at diagnosis; continuous variables, median (range); dichotomized variables (N, %). ALC: absolute lymphocyte count; AMC: absolute monocyte count; ANC: absolute neutrophil count; WBC: white blood cell count; PB: peripheral blood; BM: bone marrow. Mann-Whitney test or Kruskal-Wallis test. The bold values represent p values that are statistically significant; $p < 0.05$.

| | All patients | CMML with IMCs | CMML w/o IMCs | P value |
|---|---|---|---|---|
| ***Number*** | 580 | 351 | 229 | |
| ***Age in years; median (range)*** | 71 (18 - 95) | 70 (18 - 95) | 72 (20 - 95) | 0.06 |
| ***Males; N (%)*** | 392 (68) | 235 (67) | 157 (69) | 0.68 |
| ***Hemoglobin, g/dL; median (range)*** | 10.7 (4.3 - 17) | 10.5 (4.3 - 16.9) | 11.3 (6.4 - 17) | **0.005** |
| ***WBC x$10^9$/L; median (range)*** | 12.9 (1.3- 264.8) | 18.4 (1.8-185.7) | 7.9 (1.3- 264.8) | **<0.0001** |
| ***ANC x $10^9$/L; median (range)*** | 6.2 (0 - 151) | 8.9 (0.06 - 151) | 3.8 (0 – 142.9) | **<0.0001** |
| ***AMC x $10^9$/L; median (range)*** | 2.7 (0 - 84) | 3.3 (0.1 - 84) | 1.7 (0 – 24.6) | **<0.0001** |
| ***ALC x $10^9$/L; median (range)*** | 1.9 (0 - 22) | 2.15 (0 - 14.5) | 1.6 (0.2 - 22) | **<0.0001** |
| ***Platelets x $10^9$/L; median (range)*** | 101 (7 - 1473) | 97 (7 - 1473) | 109 (12 - 1277) | 0.17 |
| ***Abnormal cytogenetics; N (%)*** | 178/551 (32) | 114/330 (35) | 64/221 (29) | 0.16 |
| ***Proliferative MP-CMML; N (%)*** | 288 (50) | 231 (66) | 57 (25) | **<0.0001** |
| ***Dysplastic MD-CMML; N (%)*** | 292 (50) | 120 (34) | 172 (75) | |
| ***CMML-0; N (%)*** | 328 (57) | 172 (49) | 156 (68) | **<0.0001** |
| ***CMML-1; N (%)*** | 147 (25) | 103 (29) | 44 (19) | |
| ***CMML-2; N (%)*** | 104 (18) | 75 (21) | 29 (13) | |
| ***Next generation*** | 342 | 201 | 141 | |
| ***sequencing analysis; N*** | 180 (53) | 104 (52) | 76 (55) | 0.59 |
| ***TET2*** | 16 (5) | 9 (4) | 7 (5) | 0.83 |
| ***DNMT3A*** | 6 (2) | 5 (3) | 1 (1) | 0.19 |
| ***IDH1*** | 20 (6) | 12 (6) | 8 (6) | 0.9 |
| ***IDH2*** | 200 (54) | 130 (59) | 70 (46) | **0.01** |
| ***ASXL1*** | 13 (4) | 10 (5) | 3 (2) | 0.16 |
| ***EZH2*** | 33 (10) | 17 (8) | 16 (11) | 0.37 |
| ***RUNX1*** | 3 (1) | 2 (1) | 1 (1) | 0.77 |
| ***BCOR*** | 17 (5) | 7 (3) | 10 (7) | 0.13 |
| ***SF3B1*** | 158 (46) | 99 (49) | 59 (42) | 0.19 |
| ***SRSF2*** | 25 (7) | 11 (5) | 14 (10) | 0.12 |
| ***U2AF1*** | 11 (3) | 7 (3) | 4 (3) | 0.73 |
| ***ZRSR2*** | 30 (9) | 17 (8) | 13 (9) | 0.8 |
| ***JAK2 V617F*** | 1 (0.3) | 1 (0.5) | 0 (0) | 0.3 |
| ***MPL*** | 50 (15) | 33 (16) | 17 (12) | 0.25 |
| ***CBL*** | 19 (6) | 10 (5) | 9 (6) | 0.57 |
| ***KRAS*** | 51 (15) | 32 (16) | 19 (13) | 0.53 |
| ***NRAS*** | 10 (3) | 7 (3) | 3 (2) | 0.45 |
| ***PTPN11*** | 3 (1) | 3 (1.5) | 0 (0) | 0.07 |
| ***CSF3R*** | 11 (3) | 4 (2) | 7 (5) | 0.12 |
| ***C-KIT*** | 9 (3) | 6 (3) | 3 (2) | 0.62 |
| ***FLT3TKD*** | 1 (0.3) | 0 (0) | 1 (1) | 0.18 |
| ***NPM1*** | 10 (3) | 7 (3) | 3 (2) | 0.45 |
| ***Tp53*** | 40 (12) | 25 (12) | 15 (11) | 0.6 |
| ***SETBP1*** | | | | |
| ***Leukemic transformation; N (%)*** | 116 (20) | 73 (21) | 43 (19) | 0.55 |
| ***Deaths; N (%)*** | 398 (69) | 245 (70) | 153 (67) | 0.44 |

**Supplementary Table 2**. List and references of antibodies used in indicated assays.

| REAGENT | COLOR | CAT. NUMBER | CLONE | SOURCE |
|---|---|---|---|---|
| ***Antibodies for Spectral flow cytometry*** | | | | |
| CD24 | BB700 | 566525 | ML5 | BD Biosciences |
| CD1c | BUV563 | 748724 | F10/21A3 | BD Biosciences |
| CD141 | BV421 | 565321 | 1A4 | BD Biosciences |
| CD101 | BV605 | 747548 | V7.1 | BD Biosciences |
| CD163 | BUV737 | 741863 | GHI/61 | BD Biosciences |
| CD34 | BUV661 | 749902 | 581 | BD Biosciences |
| CD36 | BUV496 | 750114 | CLB-IVC7 | BD Biosciences |
| CD5 | BV750 | 747108 | UCHT2 | BD Biosciences |
| CD45RA | BUV395 | 740315 | 5H9 | BD Biosciences |
| CD15 | PE-Dazzle 594 | 562463 | HI98 | BD Biosciences |
| CD16 | BUV805 | 748850 | 3G8 | BD Biosciences |
| FcER1a | eFluor 450 | 48-5899-41 | AER-37 (CRA1) | Thermofisher |
| NKG2D | PerCP-eFluor 710 | 46-5878-41 | 1D11 | Thermofisher |
| CD3 | PerCP-eFluor 710 | 46-0037-41 | OKT3 | Thermofisher |
| CD7 | PerCP-eFluor 710 | 46-0078-41 | 4H9 | Thermofisher |
| CD19 | PerCP-eFluor 710 | 46-0198-42 | SJ25C1 | Thermofisher |
| CD123 | superbright436 | 62-1239-42 | 6H6 | Thermofisher |
| SLAN | FITC | 130-117-371 | DD1 | Miltenyibiotech |
| AXL | APC | FAB154A | #108724 | R&D |
| CD66b | Alexa Fluor 647 | 305109 | G10F5 | Biolegend |
| CD89 | APC-Fire 750 | 354115 | A59 | Biolegend |
| CCR2 | BV785 | 357233 | K036C2 | Biolegend |
| CD11b | BV650 | 101239 | M1/70 | Biolegend |
| CD33 | BV570 | 303417 | WM53 | Biolegend |
| HLA-DR | PE-Fire810 | | | Biolegend |
| CD14 | Sparkle blue 550 | 367147 | 63D3 | Biolegend |
| CD56 | PE-Fire 700 | 392427 | NCAM | Biolegend |
| CXCR4 | PE-CY5 | 306508 | 12G5 | Biolegend |
| CD45 | Percp | 304026 | HI30 | Biolegend |
| CD10 | PE-CY7 | 312214 | HI10A | Biolegend |
| CD8 | BV510 | 344732 | SK1 | Biolegend |
| CD88 | PE | 344303 | C5aR | Biolegend |
| CD47 | BV711 | 563761 | B6H12 | BD Biosciences |
| CD4 | CfluorYG584 | | SK3 | CYTEK |
| ***Antibodies for conventionnal flow cytometry*** | | | | |
| CD24 | BUV395 | 563818 | ML5 | BD Biosciences |
| CD15 | PE-Dazzle 594 | 562463 | HI98 | BD Biosciences |
| CD16 | APC-H7 | 560195 | 3G8 | BD Biosciences |
| CD3 | APC | 555342 | HIT3a | BD Biosciences |
| CD19 | APC | 55415 | SJ25C1 | BD Biosciences |
| CD66b | FITC | 555724 | G10F5 | BD Biosciences |
| CD11b | Percp-CY5.5 | 301328 | ICRF44 | Biolegend |
| CD33 | PE | 555450 | WM53 | BD Biosciences |
| HLA-DR | BV421 | 562804 | G46-6 | Biolegend |
| CD14 | PE-CY7 | A22331 | RMO52 | Beckman Coulter |
| CD56 | APC | IM2474 | N901 | Beckman Coulter |
| CD45 | BV510 | 304036 | HI30 | Biolegend |
| ***Antibodies for semi-solid culture analysis*** | | | | |
| CD163 | BUV737 | | | |
| CD15 | PE-Dazzle 594 | 562463 | HI98 | BD Biosciences |
| CD66b | Alexa Fluor 647 | 305109 | G10F5 | Biolegend |
| CD11b | BUV805 | | | Biolegend |
| CD45 | BV510 | 304036 | HI30 | Biolegend |
| CD14 | Sparkle blue 550 | 367147 | 63D3 | Biolegend |
| CD36 | BUV496 | 750114 | CLB-IVC7 | BD Biosciences |
| GPA | PE | | | |
| ***Antibodies for proliferation assays*** | | | | |
| CD3 | BUV738 | | UCHT1 | BD Biosciences |
| CD4 | APC | | OKT4 | Biolegend |
| CD8 | BV786 | | RPA-T8 | BD Biosciences |
| ***Antibodies for bone marrow CD34$^{+}$ analysis*** | | | | |
| CD45 | APC-H7 | 561863 | HI30 | BD Biosciences |
| CD34 | PE-CY7 | 560710 | | BD Biosciences |
| CD38 | eFluor 450 | 48-0388-42 | | Thermofisher |
| CXCR1 CD181 | BV650 | 749071 | | BD Biosciences |
| CXCR2 CD182 | BUV737 | 743422 | | BD Biosciences |

| | | | | |
|---|---|---|---|---|
| ***Antibodies for intracellular staining*** | | | | |
| CD19 | APC | 55415 | SJ25C1 | BD Biosciences |
| CD56 | APC | IM2474 | N901 | Beckman Coulter |
| CD66b | FITC | 555724 | G10F5 | BD Biosciences |
| CD3 | BUV737 | 612750 | UCHT1 | BD Biosciences |
| CD16 | Pacific blue | B36292 | 3G8 | Beckman Coulter |
| CD45 | BV510 | 304036 | HI30 | Biolegend |
| CD14 | BV605 | 564054 | M5E2 | BD Biosciences |
| CD15 | AF-700 | 301920 | HI98 | Biolegend |

**Supplementary Table 3.** Clinical and laboratory characteristics of CMML patients and age-matched controls tested by spectral or by conventional flow cytometry (learning cohort and validation cohorts). Continuous variables, median (range); dichotomized variables (N, %).). ALC: absolute lymphocyte count; AMC: absolute monocyte count; ANC: absolute neutrophil count; WBC: white blood cell count; GFM: groupe francophone des myelodysplasies. Mann-Whitney test or Kruskal-Wallis test. The bold values represent p values that are statistically significant; $p < 0.05$. No information available for young donors.

| | *Spectral flow cytometry* | | | *Conventional flow cytometry* | | | | |
|---|---|---|---|---|---|---|---|---|
| | *Cohort* | | | | *Learning cohort* | | *Validation cohort* | |
| | *Aged controls* | *CMML* | *p value* | *Aged controls* | *CMML* | *p value* | *CMML* | *p value* |
| *Number* | **10** | **27** | | **64** | **209** | | **160** | |
| *Age in years* | 71.5 (65-79) | 76.0 (60-92) | 0.17 | 74.0 (65-94) | 75.0 (50-93) | 0.95 | 75.0 (42-95) | 0.91 |
| *Males; N (%)* | 4 (40%) | 16 (60%) | | 28 (46%) | 132 (63%) | | 92(58%) | |
| *Hemoglobin, g/dL* | 13.9 (13.1-15.8) | 12.4 (8.4-16.2) | 0.01 | 13.4 (10.7-16.8) | 11.9 (6.4-17.2) | **<0.0001** | 12.0 (6.1-16.5) | **<0.0001** |
| *WBC x10$^9$/ L* | 5.5 (4.4-7.7) | 7.2 (3.1-19.9) | 0.085 | 6.9 (4.5-12.5) | 10.7 (2.7-141.9) | **<0.0001** | 9.55 (2.6-76) | **<0.0001** |
| *ANC x 10$^9$/L* | 3.1 (2.3-5.8) | 3.2 (1.2-9.4) | 0.64 | 4.3 (1.9-7.0) | 5.4 (0.5-65.9) | **0.0042** | 5.2 (0.1-68.8) | 0.034 |
| *Neutrophils (%)* | 57.3 (40.0-75.3) | 50.3 (29.9-77.0) | 0.129 | 61.5 (36.0-78.0) | 51.0 (7.0-93.8) | **<0.0001** | 52.7 (2.6-92) | 0.0002 |
| *AMC x 10$^9$/L* | 0.4 (0.3-0.6) | 1.65 (1-6.6) | <0.0001 | 0.6 (0.3-0.9) | 2.22 (1-59.7) | **<0.0001** | 1.85 (0.5-28.8) | **<0.0001** |
| *Monocytes (%)* | 7.5 (6.0-11.0) | 28 (12.8-42.0) | <0.0001 | 8.7 (3.7-17.8) | 23.0 (10.0-64.0) | **<0.0001** | 21.4 (10.0-57.7) | **<0.0001** |
| *ALC x 10$^9$/L* | 1.5 (1.1-3.1) | 1.4 (0.4-3.8) | 0.27 | 1.9 (1.2-2.9) | 1.9 (0.3-10) | 0.6 | 1.7 (0.4-12) | 0.4 |
| *Lymphocytes (%)* | 29.0 (15.6-43.8) | 22.2 (3.1-39.5) | 0.006 | 27 (16-51) | 18 (1.4-50) | **<0.0001** | 19 (1-63) | **<0.0001** |
| *Platelets x 10$^9$/L* | 227 (183-369) | 127 (25-1097) | 0.0009 | 234 (152-431) | 111 (12-688) | **<0.0001** | 140 (13-1090) | **<0.0001** |
| *Abnormal karyotype/tested; N (%)* | NA | 2 / 18 (11%) | | NA | 46 / 190 (24%) | | 22 / 104 (21%) | |
| *Proliferative; N (%)* | NA | 23 (85%) | | NA | 83 (40%) | | 48 (34%) | |
| *Dysplastic; N (%)* | NA | 4 (15%) | | NA | 126 (60%) | | 94 (66%) | |
| *CMML-0; N (%)* | NA | 15 (55%) | | NA | 104 (49%) | | 62 (43%) | |
| *CMML-1; N (%)* | NA | 10 (38%) | | NA | 77 (36.5%) | | 64 (44%) | |
| *CMML-2; N (%)* | NA | 2 (7%) | | NA | 31 (14.5%) | | 19 (13.0%) | |
| *Classical monocytes. (% total monocytes)* | NA | 97.5% (94.4-99.5) | | NA | 96.9%. (69-99.5) | | 96.7%. (54-99.6) | |
| *GFM prognosis model, N* | | 27 CMML tested | | | 180 CMML tested | | 140 CMML tested | |
| *GFM, low; N (%)* | NA | 21 (78%) | | NA | 93 (52%) | | 107 (76%) | |
| *GFM, intermed; N (%)* | NA | 4 (15%) | | NA | 59 (33%) | | 23 (17%) | |
| *GFM, high; N (%)* | NA | 2 (7%) | | NA | 28 (15%) | | 10 (7%) | |
| *NGS analysis; N* | | 27 CMML tested | | | 180 CMML tested | | 121 CMML tested | |
| *TET2 mutated N (%)* | NA | 23 (85%) | | NA | 120 (67%) | | 89 (73%) | |
| *SRSF2 mutated N (%)* | NA | 15 (55%) | | NA | 64 (36%) | | 45 (37%) | |
| *ASXL1 mutated N (%)* | NA | 11 (41%) | | NA | 48 (27%) | | 44 (36%) | |
| *NRAS mutated N (%)* | NA | 2 (7%) | | NA | 22 (12%) | | 13 (11%) | |
| *KRAS mutated N (%)* | NA | 3 (11%) | | NA | 21 (12%) | | 16 (13%) | |
| *CBL mutated N (%)* | NA | 5 (19%) | | NA | 14 (8%) | | 14 (11%) | |
| *AML1 mutated N (%)* | NA | 2 (7%) | | NA | 33 (18%) | | 17 (14%) | |

**Supplementary Table 4.** Correlation between iGRANs (x10$^9$/L) and clinical and biological disease parameters. Mann-Whitney test or Kruskal-Wallis test.

| ***Parameters*** | **Number** | **Median [95%CI]** | **Significance** |
|---|---|---|---|
| ***A - Categorical variables*** | | | |
| ***Sex*** | | | NS |
| *Females* | 77 | 0.2 [0.05-2.06] | |
| *Males* | 132 | 0.22 [0.04-2.05] | |
| ***WHO CMML subtype 2016*** | | | **0.019** |
| *CMML-0* | 101 | 0.15 [0.04-0.76] | |
| *CMML-1* | 77 | 0.28 [0.06-2.58] | |
| *CMML-2* | 31 | 0.91 [0.07-5.79] | |
| ***FAB CMML subtype*** | | | **P<0001** |
| *Dysplastic* | 126 | 0.07 [0.03-0.21] | |
| *Proliferative* | 83 | 2.23 [0.62-7.54] | |
| ***ASXL1 mutation*** | | | **P<0001** |
| *Absent* | 150 | 0.13 [0.04-0.71] | |
| *Present* | 53 | 2.49 [0.28-5.64] | |
| ***TET2 mutation*** | | | NS |
| *Absent* | 64 | 0.18 [0.04-3.42] | |
| *Present* | 140 | 0.21 [0.05-1.45] | |
| ***SRSF2 mutation*** | | | NS |
| *Absent* | 121 | 0.17 [0.04-1.07] | |
| *Present* | 73 | 0.36 [0.07-2.88] | |
| ***AML1 mutation*** | | | NS |
| *Absent* | 168 | 0.19 [0.04-1.75] | |
| *Present* | 35 | 0.38 [0.07-5.01] | |
| ***NRAS mutation*** | | | **0.005** |
| *Absent* | 179 | 0.17 [0.04-1.51] | |
| *Present* | 24 | 1.31 [0.25-5.57] | |
| ***KRAS mutation*** | | | NS |
| *Absent* | 177 | 0.23 [0.05-2.15] | |
| *Present* | 26 | 0.07 [0.03-0.58] | |
| ***CBL mutation*** | | | NS |
| *Absent* | 187 | 0.19 [0.04-1.72] | |
| *Present* | 16 | 1.52 [0.1-4.65] | |
| ***GFM prognostic score*** | | | **P<0001** |
| *Low* | 116 | 0.07 [0.03-0.21] | |
| *Int* | 62 | 1.12 [0.21-3.03] | |
| *High* | 30 | 5.32 [2.35-12.06] | |
| ***CPSS risk stratification*** | | | **P<0001** |
| *Low* | 75 | 0.07 [0.03-0.19] | |
| *Int-1* | 55 | 0.58 [0.10-2.15] | |
| *Int-2* | 56 | 1.19 [0.24-9.45] | |
| *High* | 5 | 2.19 [1.12-5.79] | |
| ***CPSS-M risk stratification*** | | | **P<0001** |
| *Low* | 34 | 0.07 [0.03-0.19] | |
| *Int-1* | 46 | 0.08 [0.04-0.25] | |
| *Int-2* | 51 | 0.61 [0.1-2.75] | |
| *High* | 49 | 2.49 [0.34-7.55] | |
| ***B - Continuous variables*** | **Number** | **Spearman correlation** | **Significance** |
| ***Age*** | 209 | -0.05 | NS |
| ***Hemoglobin, g/dL*** | 209 | -0.3 | **P<0001** |
| ***Platelets, x10$^9$/L*** | 209 | -0.15 | **0.02** |
| ***WBC count, x10$^9$/L*** | 209 | 0.71 | **P<0001** |
| ***Neutrophils (%)*** | 209 | -0.0093 | NS |
| ***Monocytes (%)*** | 209 | -0.03 | NS |
| ***Lymphocytes (%)*** | 208 | -0.47 | **P<0001** |
| ***IMC (%)*** | 187 | 0.73 | **P<0001** |

**Supplementary Table 5.** Correlation between iGRANs (expressed in fraction or absolute number x $10^9$/L) and overall survival or event-free survival (EFS, defined as the time between diagnosis and AML transformation, death, or last follow-up) using Cox models, Multivariate model were built after adjusting for potential confounders (Age, WBC > 15x $10^9$/L, hemoglobin <10g/L, platelets < 100 x $10^9$/L, *ASXL1* mutations). The proportional hazard hypothesis was verified by visual display of the cumulative sums of martingale residuals.

| **Analysis with continuous data** | **Hazard ratio [95%CI] (univariate model)** | **P – value** | **Hazard ratio [95%CI] (multivariate model)** | **P – value** |
|---|---|---|---|---|
| ***Overall survival (OS)*** | | | | |
| ***iGRAN (%)*** | 1.029 [1.019-1.038] | <0.0001 | 1.014 [1.00-1.026] | 0.0115 |
| ***iGRAN ($10^9$/L)*** | 1.104 [1.074-1.135] | <0.0001 | 1.059 [1.02-1.099] | 0.0028 |
| ***Event-free survival (EFS)*** | | | | |
| ***iGRAN (%)*** | 1.028 [1.019-1.037] | <0.0001 | 1.016 [1.00-1.026] | 0.0049 |
| ***iGRAN ($10^9$/L)*** | 1.103 [1.074-1.132] | <0.0001 | 1.067 [1.03-1.105] | 0.0003 |
| **Analysis with an iGRAN cutpoint** | **Hazard ratio [95%CI] (univariate model)** | **P – value** | **Hazard ratio [95%CI] (multivariate model)** | **P – value** |
| ***Overall survival (OS)*** | | | | |
| ***iGRAN (14%)*** | 5.93 [3.364-10.454] | <0.0001 | 3.678 [1.850-7.305] | 0.0002 |
| ***iGRAN (0.4x$10^9$/L)*** | 5.49 [3.183-9.470] | <0.0001 | 3.115 [1.482-6.548] | 0.0027 |
| ***Event-free survival (EFS)*** | | | | |
| ***iGRAN (14%)*** | 5.10 [2.993-8.684] | <0.0001 | 3.003 [1.592-5.663] | 0.0007 |
| ***iGRAN (0.4x$10^9$/L)*** | 4.79 [2.857-8.032] | <0.0001 | 2.579 [1.292-5.149] | 0.0003 |

**Supplementary Table 6.** Variant allele frequency (VAF) detected by Whole-exome sequencing (WES) performed in monocytes, T cells and iGRANs sorted from 14 CMML blood samples. All mutations were found in both iGRANs and monocytes, but not or at low VAF in T lymphocytes.

| | | Variant allele frequency | | |
|---|---|---|---|---|
| | **Mutated gene** | **T Lymphocytes** | **Monocytes** | **iGRANs** |
| ***WES4*** | *ROCK2* | 2.83% | 58.4% | 50% |
| | *TMEM247* | 9.44% | 23.44% | 23% |
| | *FILIP1* | 5.14% | 50% | 43.88% |
| | *COL1A2* | 8.2% | 54.55% | 48.39% |
| | *FCN2* | 4.67% | 42.65% | 44% |
| | *KRAS* | 4.46% | 36.88% | 33.76% |
| | *MGAT4C* | 9.68% | 44.19% | 35.59% |
| | *EPB42* | 6.43% | 51.27% | 48.6% |
| | *GMEB2* | 6.58% | 49.46% | 50.36% |
| | *C6orf223* | 6.67% | 42.22% | 32,07% |
| ***WES10*** | *NSD1* | 0.61% | 20.43% | 26.97% |
| | *GIMAP1* | 1.55% | 19.8% | 33.16% |
| | *RIOX1* | 0% | 14.81% | 21.95% |
| | *ATXN3* | 9.09% | 42.25% | 27.96% |
| | *TULP2* | 0% | 13.56% | 26.12% |
| | *TET2* | 14.76% | 27.94% | 28.11% |
| | *NRAS* | 3% | 5% | 31% |
| | *PTPN11* | 5% | 8% | 10% |
| ***WES8*** | *IFNG* | 3.9% | 50.9% | 86% |
| | *MGAT4D* | 15.8% | 80% | 39% |
| | *DDX20* | 0% | 34.4% | 25.9% |
| | *TET2* | 26.5% | 43.7% | 49.2% |
| | *TET2* | 57.1% | 55.7% | 50% |
| | *NRAS* | 23.2% | 35% | 42% |
| | *ASXL1* | 28.4% | 47.8% | 41% |
| | *PIR* | 35% | 54% | 49% |
| | *GRM4* | 18.75% | 49% | 38% |
| | *ZPLD1* | 28.6% | 47% | 46% |
| | *LAMB2* | 26.2% | 51% | 54% |
| ***WES3*** | *ZRANB2* | 1.65% | 32.61% | 14.57% |
| | *NRAS* | 4.75% | 16.98% | 27.55% |
| | *NBEAL1* | 8.86% | 42.86% | 40% |
| | *MAST4* | 3.15% | 51.92% | 40.72% |
| | *TNFRSF10A* | 6.36% | 49.3% | 43.88% |
| | *C8orf34* | 6.97% | 43.48% | 38.06% |
| | *ZFHX4* | 0% | 50% | 36% |
| | *KIAA2026* | 8.22% | 45.1% | 37.5% |
| | *PRSS3* | 7.68% | 18.09% | 16.39% |
| | *KRAS* | 0.46% | 25.42% | 12.9% |
| | *PTPN11* | 6.47% | 48.54% | 47.62% |
| | *LDHD* | 1.35% | 28.18% | 12.73% |
| | *SRSF2* | 8.08% | 42.31% | 47.97% |
| | *APLP1* | 8.06% | 48.39% | 42.31% |
| | *ATP1A3* | 6.67% | 48.85% | 48.08% |
| | *TET2* | 25% | 51% | 54% |

| | | | | |
|---|---|---|---|---|
| **WES5** | *ADAMTS20* | 20.6% | 45.8% | 49.4% |
| | *ACCS* | 19.7% | 51.9% | 41.4% |
| | *TUSC1* | 15.8% | 56.1% | 57.9% |
| | *COL5A3* | 20.8% | 50% | 51.9% |
| | *APPL1* | 26.4% | 58.1% | 46.4% |
| | *TET2* | 12.2% | 48.3% | 46.1% |
| | *TET2* | 20.8% | 40.4% | 46.4% |
| | *SRSF2* | 26.22% | 48.2% | 42.3% |
| | *MRGPRE* | 11.5% | 40.5% | 45.4% |
| | *ADAMTSL2* | 10% | 60% | 40.2% |
| | *E2F3* | 19.6% | 44.8% | 46.7% |
| | *MEGF6* | 28.3% | 35% | 51.2% |
| | *ASXL1* | 20% | 23% | 23.8% |
| | *NF1* | 10.8% | 48.7% | 28% |
| | *CBL* | 42% | 98% | 99% |
| | *FOS* | 27.8% | 48.5% | 43% |
| | *UPF3A* | 20% | 40.5% | 36.8% |
| **WES2** | *C1orf185* | 3.03% | 38.46% | 46.43% |
| | *NRAS* | 8.66% | 47.46% | 39.39% |
| | *FMN2* | 4.45% | 11.11% | 11.11% |
| | *ARSI* | 8.74% | 49.74% | 47.21% |
| | *PSMB5* | 9.26% | 42.02% | 47.54% |
| | *DNAH9* | 4.71% | 24.36% | 20.5% |
| | *NF1* | 7.02% | 29.23% | 18.78% |
| | *SETBP1* | 7.31% | 46.39% | 37.57% |
| | *ASXL1* | 9.52% | 27.66% | 27.19% |
| **WES1** | *MPL* | 2.35% | 10.76% | 25.25% |
| | *KANK4* | 7.69% | 60.49% | 48.19% |
| | *NRAS* | 4.69% | 47.18% | 50.37% |
| | *NPR1* | 8.87% | 60.33% | 59.9% |
| | *CFAP221* | 4.2% | 47.54% | 48.35% |
| | *TET2* | 9.75% | 51.25% | 57.33% |
| | *TET2* | 7.11% | 46.63% | 49.16% |
| | *ZNF318* | 6.08% | 41.03% | 50.83% |
| | *RIMS1* | 2.8% | 37.96% | 22.02% |
| | *CFTR* | 4.76% | 41.46% | 46.67% |
| | *ANK1* | 8% | 57.42% | 51.85% |
| | *RSPO2* | 4.14% | 52.14% | 43.97% |
| | *ST14* | 8.62% | 47.55% | 46.74% |
| | *FOXO1* | 6.74% | 64.1% | 37.84% |
| | *FSCB* | 0% | 38.89% | 43.33% |
| | *TRIP11* | 1.4% | 20.31% | 13.21% |
| | *ALDH1A2* | 7.07% | 52.1% | 54.33% |
| | *CDH13* | 2.41% | 39.44% | 25.09% |
| | *SRSF2* | 6.59% | 45.74% | 49.56% |
| | *DOHH* | 1.08% | 34.15% | 27.12% |
| | *CENPB* | 4.97% | 49.66% | 40.48% |
| | *EMILIN3* | 7% | 43.33% | 58.7% |
| **WES9** | *FSIP2* | 3.77% | 46.6% | 52.38% |
| | *CCDC80* | 1.7% | 16.33% | 23.53% |

| | | | | |
|---|---|---|---|---|
| | *ILDR1* | 5.97% | 33.33% | 50% |
| | *TET2* | 5.45% | 39.02% | 48.54% |
| | *TPPP* | 6.49% | 54.22% | 69.41% |
| | *PCDHGA1* | 7.77% | 48.24% | 42.86% |
| | *DNAH8* | 7.25% | 45.45% | 46.46% |
| | *TNFAIP3* | 8.62% | 48.48% | 38.64% |
| | *CHRNA6* | 1.23% | 16.2% | 10.62% |
| | *C10orf71* | 7.61% | 51.9% | 38.46% |
| | *OR9G1;OR9G9* | 4.27% | 18.89% | 29.89% |
| | *AHNAK* | 7.41% | 46.49% | 41.77% |
| | *CABP2* | 8.23% | 45.59% | 42.05% |
| | *ETV6* | 5.75% | 49.64% | 44.72% |
| | *PDZRN4* | 4.55% | 47.37% | 45.21% |
| | *PIWIL1* | 5.83% | 53.57% | 46.67% |
| | *LMO7* | 2.22% | 50.59% | 49% |
| | *SETD1A* | 7.27% | 43.97% | 40.38% |
| | *PRDM7* | 9.54% | 41.36% | 52.34% |
| | *LOC100506388* | 4.26% | 36.59% | 42.86% |
| | *HNF1B* | 4.98% | 44.77% | 47.65% |
| | *SRSF2* | 5.13% | 43.24% | 49.02% |
| | *TSHZ1* | 0.42% | 15.74% | 13.09% |
| | *ASXL1* | 8.15% | 46.58% | 47.18% |
| | *C1QTNF6* | 5.83% | 54.4% | 54.48% |
| | *TUBGCP6* | 4.51% | 50.28% | 32.23% |
| | *KRAS* | 0.51% | 7% | 11.11% |
| | *RIT1* | 2.56% | 28.87% | 14% |
| ***WES16*** | *PRAMEF12* | 3.45% | 55% | 50% |
| | *SLC25A34* | 1.1% | 46.88% | 52.94% |
| | *MPL* | 1.92% | 45.1% | 44.04% |
| | *ERICH3* | 8.75% | 51.35% | 52.73% |
| | *VAV3* | 5.32% | 45.78% | 46.21% |
| | *NRAS* | 3.64% | 40.74% | 50% |
| | *CACNA1S* | 3.7% | 44.72% | 46.12% |
| | *TET2* | 7.37% | 40.28% | 46.51% |
| | *FAT4* | 2.17% | 39.37% | 47.49% |
| | *MND1* | 8.7% | 50% | 51.43% |
| | *CACNA1B* | 7.32% | 51.85% | 55.77% |
| | *IRAK4* | 9.09% | 42.31% | 44.68% |
| | *SRCAP* | 4.68% | 48.53% | 46.83% |
| | *ASXL1* | 3.57% | 36.84% | 42.99% |
| ***WES11*** | *PRAMEF12* | 5.56% | 48.15% | 41.94% |
| | *INO80B* | 4.69% | 38.64% | 56% |
| | *UNC80* | 8.33% | 42.91% | 41.46% |
| | *NEK4* | 9.49% | 50% | 51.46% |
| | *MARCH11* | 8.6% | 43.09% | 32.53% |
| | *NLRP10* | 8.97% | 55.68% | 56.14% |
| | *NLRX1* | 7.6% | 54.22% | 50% |
| | *UNC79* | 6.06% | 38.17% | 44.51% |
| | *MGA* | 5.63% | 43.14% | 34.15% |
| | *CDH19* | 6.94% | 50.62% | 50% |
| | *SRSF2* | 12% | 52% | 66% |

| | | | | |
|---|---|---|---|---|
| | *ASXL1* | 9.67% | 29.16% | 28.20% |
| **WES14** | *NRAS* | 5.56% | 42.19% | 53.85% |
| | *ACMSD* | 6.45% | 46.51% | 42.05% |
| | *SP140* | 3.45% | 44.66% | 40.62% |
| | *UGT1A8* | 9.3% | 62.26% | 43.14% |
| | *MUC4* | 0.39% | 16.36% | 17.51% |
| | *PRSS3* | 7.21% | 23.86% | 25.47% |
| | *UBN1* | 8.57% | 53.57% | 41.18% |
| | *KCNC3* | 4.49% | 44.24% | 42.95% |
| | *ASXL1* | 5.45% | 45% | 34.83% |
| | *TET2* | 15% | 59% | 46% |
| | *TET2* | 19% | 46.15% | 49.09% |
| | *SRSF2* | 14% | 52% | 53% |
| **WES13** | *USH2A* | 0.91% | 45.68% | 28.93% |
| | *VAX2* | 8.39% | 47.06% | 45.26% |
| | *TET2* | 9.43% | 45.83% | 34.85% |
| | *TET2* | 5% | 47.62% | 40.74% |
| | *SMAD1* | 3.75% | 46.15% | 50.57% |
| | *MUC22* | 2.29% | 14.64% | 11.19% |
| | *DPP3* | 0% | 36.36% | 42.86% |
| | *CBL* | 4.95% | 82% | 79.07% |
| | *SCNN1G* | 4.35% | 52% | 51.18% |
| | *KRT16* | 4.76% | 36.21% | 31.08% |
| | *TMC8* | 2.78% | 42.22% | 56.52% |
| | *RRBP1* | 3.57% | 22.58% | 21.33% |
| **WES15** | *RPTN* | 0.58% | 43.67% | 50.99% |
| | *CTNNA2* | 4.67% | 43.59% | 48.06% |
| | *ERFE* | 3.23% | 68% | 50.98% |
| | *MUC4* | 0.26% | 12.86% | 12.25% |
| | *EYS* | 1.43% | 41.77% | 46.43% |
| | *RADIL* | 6.98% | 36.21% | 47.54% |
| | *EZH2* | 6.67% | 43.3% | 39.29% |
| | *SLC46A2* | 5.19% | 55.05% | 50.39% |
| | *ARHGAP11A* | 2.56% | 45.83% | 44.44% |
| | *ALOX15* | 5.5% | 51.45% | 55.96% |
| | *DNAH9* | 6.78% | 40% | 56.25% |
| | *SRSF2* | 8.24% | 41.44% | 39.47% |
| | *ASXL1* | 5.41% | 45.71% | 39.09% |
| | *USH2A* | 0% | 38% | 62.5% |
| | *TTC39B* | 9.3% | 43.33% | 22% |
| **WES12** | *CSF3R* | 5% | 88.46% | 100% |
| | *CSF3R* | 1.61% | 30.22% | 37.41% |
| | *NPAS2* | 1.69% | 41.27% | 36.11% |
| | *LCT* | 3.03% | 56% | 45.95% |
| | *GATB* | 0% | 37.97% | 39.58% |
| | *ASB5* | 3.57% | 41.25% | 47.06% |
| | *ZDHHC11* | 0% | 17.21% | 25% |
| | *RXRB* | 3.17% | 44.44% | 50.99% |
| | *CDK14* | 1.1% | 32.41% | 37.66% |
| | *CUX1* | 0% | 48.75% | 56.52% |
| | *DEFB105A;DEFB105B* | 2.04% | 29.82% | 34.29% |

| | | | | |
|---|---|---|---|---|
| | *DEFB105A;DEFB105B* | 0.67% | 28.14% | 32.76% |
| | *FGF20* | 0.76% | 33.33% | 39.29% |
| | *MMP12* | 0% | 83.97% | 91.84% |
| | *DAZAP2* | 6.14% | 38.01% | 30.34% |
| | *LRP1* | 0.85% | 24.09% | 32.23% |
| | *C15orf39* | 2% | 34.62% | 54.17% |
| | *ACSF3* | 0.8% | 38.8% | 47.56% |
| | *KRT28* | 3.45% | 49.17% | 50.62% |
| | *KRT34* | 2.7% | 38.58% | 41.05% |
| | *SRSF2* | 5.71% | 38.64% | 34.48% |
| | *UBA2* | 0.83% | 10.12% | 12.5% |
| | *USP29* | 2.33% | 39.62% | 40% |
| | *ASXL1* | 2.17% | 32.65% | 54.29% |
| | *NBPF3* | 3.6% | 19.4% | 12% |
| | *ALG14* | 0% | 10.79% | 13% |
| | *UGDH* | 4.74% | 13% | 16.03% |
| | *MAP1B* | 1.52% | 5% | 11.97% |
| | *ZSWIM8* | 7.34% | 15% | 23.62% |

**Supplemental table 7.** Estimate fixation time (years) in function of relative fitness, derived from a Moran birth-death process. Fixation times have been derived from calculated CMML relative fitness (r, from 1.08 to 5, N=10), in a constant size population of 100,000 stem cells, with different division rates per year (d, from 0.5 to 3, N=6) without (w/o) CXCL8 and with CXCL8. The decrease in fixation time between the condition with CXCL8 and without CXCL8 is calculated (%).

| CMML patients | d | Relative fitness (r) | | Fixation time (years) | | Decrease in fixation time with CXCL8 compared to w/o CXCL8 (%) |
|---|---|---|---|---|---|---|
| | | w/o CXCL8 | with CXCL8 | w/o CXCL8 | with CXCL8 | |
| CMML 2574 | 0.5 | 1.16 | 1.49 | 310.8 | 117.0 | 62.4 |
| CMML 2812 | 0.5 | 1.67 | 2.18 | 91.8 | 62.1 | 32.4 |
| CMML 4137 | 0.5 | 1.45 | 1.75 | 125.4 | 84.4 | 32.7 |
| CMML 3073 | 0.5 | 1.3 | 1.6 | 176.5 | 99.8 | 43.5 |
| CMML 2239 | 0.5 | 1.53 | 2.09 | 109.9 | 65.3 | 40.6 |
| CMML 2300 | 0.5 | 1.52 | 2.02 | 111.6 | 68.2 | 38.9 |
| CMML 1565 | 0.5 | 1.24 | 1.58 | 214.9 | 102.4 | 52.3 |
| CMML 1745 | 0.5 | 1.07 | 1.44 | 680.9 | 127.7 | 81.2 |
| CMML 2187 | 0.5 | 1.21 | 1.83 | 242.3 | 78.5 | 67.6 |
| CMML 2415 | 0.5 | 1.08 | 1.66 | 598.7 | 92.8 | 84.5 |
| CMML 2574 | 1 | 1.16 | 1.49 | 155.4 | 58.5 | 62.4 |
| CMML 2812 | 1 | 1.67 | 2.18 | 45.9 | 31 | 32.4 |
| CMML 4137 | 1 | 1.45 | 1.75 | 62.7 | 42.2 | 32.7 |
| CMML 3073 | 1 | 1.3 | 1.6 | 88.3 | 49.9 | 43.5 |
| CMML 2239 | 1 | 1.53 | 2.09 | 55 | 32.6 | 40.6 |
| CMML 2300 | 1 | 1.52 | 2.02 | 55.8 | 34.1 | 38.9 |
| CMML 1565 | 1 | 1.24 | 1.58 | 107.5 | 51.2 | 52.3 |
| CMML 1745 | 1 | 1.07 | 1.44 | 340.5 | 63.8 | 81.2 |
| CMML 2187 | 1 | 1.21 | 1.83 | 121.2 | 39.3 | 67.6 |
| CMML 2415 | 1 | 1.08 | 1.66 | 299.3 | 46.4 | 84.5 |
| CMML 2574 | 1.5 | 1.16 | 1.49 | 103.6 | 39 | 62.4 |
| CMML 2812 | 1.5 | 1.67 | 2.18 | 30.6 | 20.7 | 32.4 |
| CMML 4137 | 1.5 | 1.45 | 1.75 | 41.8 | 28.1 | 32.7 |
| CMML 3073 | 1.5 | 1.3 | 1.6 | 58.8 | 33.3 | 43.5 |
| CMML 2239 | 1.5 | 1.53 | 2.09 | 36.6 | 21.8 | 40.6 |
| CMML 2300 | 1.5 | 1.52 | 2.02 | 37.2 | 22.7 | 38.9 |
| CMML 1565 | 1.5 | 1.24 | 1.58 | 71.6 | 34.1 | 52.3 |
| CMML 1745 | 1.5 | 1.07 | 1.44 | 227 | 42.6 | 81.2 |
| CMML 2187 | 1.5 | 1.21 | 1.83 | 80.8 | 26.2 | 67.6 |
| CMML 2415 | 1.5 | 1.08 | 1.66 | 199.6 | 30.9 | 84.5 |
| CMML 2574 | 2 | 1.16 | 1.49 | 77.7 | 29.3 | 62.4 |
| CMML 2812 | 2 | 1.67 | 2.18 | 22.9 | 15.5 | 32.4 |
| CMML 4137 | 2 | 1.45 | 1.75 | 31.3 | 21.1 | 32.7 |
| CMML 3073 | 2 | 1.3 | 1.6 | 44.1 | 24.9 | 43.5 |
| CMML 2239 | 2 | 1.53 | 2.09 | 27.5 | 16.3 | 40.6 |
| CMML 2300 | 2 | 1.52 | 2.02 | 27.9 | 17 | 38.9 |
| CMML 1565 | 2 | 1.24 | 1.58 | 53.7 | 25.6 | 52.3 |
| CMML 1745 | 2 | 1.07 | 1.44 | 170.2 | 31.9 | 81.2 |
| CMML 2187 | 2 | 1.21 | 1.83 | 60.6 | 19.6 | 67.6 |
| CMML 2415 | 2 | 1.08 | 1.66 | 149.7 | 23.2 | 84.5 |
| CMML 2574 | 2.5 | 1.16 | 1.49 | 62.2 | 23.4 | 62.4 |
| CMML 2812 | 2.5 | 1.67 | 2.18 | 18.4 | 12.4 | 32.4 |
| CMML 4137 | 2.5 | 1.45 | 1.75 | 25.1 | 16.9 | 32.7 |
| CMML 3073 | 2.5 | 1.3 | 1.6 | 35.3 | 20 | 43.5 |
| CMML 2239 | 2.5 | 1.53 | 2.09 | 22 | 13.1 | 40.6 |
| CMML 2300 | 2.5 | 1.52 | 2.02 | 22.3 | 13.6 | 38.9 |
| CMML 1565 | 2.5 | 1.24 | 1.58 | 43 | 20.5 | 52.3 |
| CMML 1745 | 2.5 | 1.07 | 1.44 | 136.2 | 25.5 | 81.2 |
| CMML 2187 | 2.5 | 1.21 | 1.83 | 48.5 | 15.7 | 67.6 |
| CMML 2415 | 2.5 | 1.08 | 1.66 | 119.7 | 18.6 | 84.5 |
| CMML 2574 | 3 | 1.16 | 1.49 | 51.8 | 19.5 | 62.4 |
| CMML 2812 | 3 | 1.67 | 2.18 | 15.3 | 10.3 | 32.4 |
| CMML 4137 | 3 | 1.45 | 1.75 | 20.9 | 14.1 | 32.7 |
| CMML 3073 | 3 | 1.3 | 1.6 | 29.4 | 16.6 | 43.5 |
| CMML 2239 | 3 | 1.53 | 2.09 | 18.3 | 10.9 | 40.6 |
| CMML 2300 | 3 | 1.52 | 2.02 | 18.6 | 11.4 | 38.9 |
| CMML 1565 | 3 | 1.24 | 1.58 | 35.8 | 17.1 | 52.3 |
| CMML 1745 | 3 | 1.07 | 1.44 | 113.5 | 21.3 | 81.2 |
| CMML 2187 | 3 | 1.21 | 1.83 | 40.4 | 13.1 | 67.6 |
| CMML 2415 | 3 | 1.08 | 1.66 | 99.8 | 15.5 | 84.5 |
